# Alternative Splicing And Global Transcriptome Changes Associated With LPS Stimulation In Human Peripheral Blood Mononuclear Cells

**DOI:** 10.1101/2025.11.09.687437

**Authors:** Esther Chavez-Iglesias, Anatol Sucher, Nidhi Thati, Julia Trudeau, Sue Yom, Marina Sirota, Adam Olshen, Nam Woo Cho, Kord M. Kober

## Abstract

**Introduction:** Lipopolysaccharide (LPS), a major component of gram-negative bacterial cell walls, elicits strong innate immune activation and is a widely used model for studying inflammatory responses. While the transcriptional response to LPS stimulation has been characterized, the role of alternative splicing (AS) in modulating this response remains largely unexplored.

**Methods:** Using deep RNA sequencing of Peripheral Blood Mononuclear Cells from three healthy female donors, we evaluated transcriptome-wide differential gene expression and alternative splicing in response to LPS stimulation.

**Results:** Our global differential gene expression and pathway impact analyses identified 490 differentially expressed genes and 46 significantly perturbed KEGG pathways, recapitulating known LPS-induced inflammatory responses and identifying two novel signaling pathways, (e.g., SNARE interactions in vesicular transport and the mRNA surveillance pathways). Differential alternative splicing analysis revealed critical impacts on immune-related pathways, including Toll-like receptor signaling, PI3K/AKT signaling, and pro-inflammatory macrophage polarization. Notably, we identified alternative splicing events in genes such as *MyD88* and *TLR4*, which play key roles in terminating inflammatory signaling, as well as splicing of long non-coding RNAs (e.g., *MALAT1*, *PVT1*) with potential regulatory functions in immune responses.

**Discussion:** This study is the first transcriptome-wide characterization of alternative splicing in response to LPS stimulation in PBMCs. Our findings suggest that alternative splicing is a fundamental regulatory mechanism in the inflammatory response and provides potential targets for therapeutic intervention in immune-related conditions.

## 1 Introduction

Lipopolysaccharide (LPS) is a commonly used stimulant that induces an innate immune response. It is a major component of the cell wall of gram-negative bacteria and is highly immunogenic, making LPS a valuable model for studying inflammatory responses.(1) LPS stimulation is a model for bacterial infection, including sepsis. Understanding the transcriptomic response to LPS stimulation could provide a mechanistic understanding and facilitate the identification of therapeutic targets.

Alternative splicing is essential to gene expression and cell function,(2) and is a key layer of gene regulation during disease progression and stress response.(3) Alternative splicing is a common and highly conserved method to remove introns and produce functional mRNAs and has been identified to play a role in the innate immune response(4) and the response to LPS stimulation.(5–7) Alternative splicing plays a regulatory role in the termination of the Toll-like receptor signaling pathway, whereby alternative splicing of genes in the pathway produce proteins that act as negative inhibitors.(5) However, prior studies have not looked into this important mechanism transcriptome-wide.

Peripheral blood represents a common pathway for immune cell trafficking to target organs as part of the immune response.(8) Peripheral blood mononuclear cells (PBMCs), which include lymphocytes and monocytes, are essential for immune responses.(9) While the transcriptomic response to LPS has been previously characterized in other studies in numerous tissues(10) and model systems, these studies have focused primarily on specific target genes. Of the four studies identified that characterized the general transcriptomic response in healthy PBMCs, two evaluated RNA using microarrays, one used bulk RNA-seq, and one used single-cell RNA-seq. Although these studies provide insight into changes in the transcriptome associated with LPS, they are either limited to pre-defined probes, limited in coverage of less expressed genes, or in vivo evaluation limited to poly-A mRNA in CD14 monocytes. Similarly, previous studies that have evaluated for alternative splicing in response to LPS stimulation in humans have primarily focused on specific genes (e.g., MyD88) or pathways (e.g., the Toll-Like Receptor (TLR) signaling pathway).(5) In pre-clinical models, one study evaluated alternative splicing using bulk RNA-seq in the ipsilateral hippocampus in ischemic and LPS-infused rats.(10) However, this study was limited by unpaired reads, short read length (50 bp), and low coverage (∼30M). In addition to TLRs, long non-coding RNAs (lncRNA) play an important role in regulating the human innate immune response and are differentially expressed in response to bacterial LPS.(11) In terms of precision medicine, an increasing number of studies have identified sex-specific immune responses. Notably, Toll-like receptors (TLRs) have been implicated in driving these sex-specific immune responses.(12) However, the relative role of hormonal or X chromosome response is unknown. (13)

Given that the role of alternative splicing in the immune response is largely unexplored, understanding the role of alternative splicing in the immune response can improve our understanding of gene regulatory processes and identify novel therapeutic targets. The purpose of this study, using deep sequencing, is to evaluate for global differential gene expression and alternative splicing associated with LPS treatment in human PBMCs.

## 2 Materials and Methods

### 2.1 Donor peripheral blood mononuclear cells

PBMCs of three healthy adult women were purchased from Stem Cell Technologies. All women were between the ages of 48-54 and self-reported ethnicity of either Asian, Caucasian, or Black (Supplemental Table 1). PBMCs had >98% viability at isolation with the vendor and were stored in liquid nitrogen upon arrival.

### 2.2 PBMC treatment with lipopolysaccharide

Donor samples (1×10^8^ cells) were thawed in a water bath set to 37°C and subsequently washed twice with pre-warmed complete RPMI 1640 medium (STEMCELL Technologies, Catalog 36750), which was supplemented with GlutaMAX (Fisher Scientific, Catalog A4192301) and 5% FBS (Fisher Scientific, Catalog A5669401). Cell viability and cell counts were measured immediately following the initial thaw and again after the final wash using the Countess 3 FL (Fisher Scientific, Catalog AMQAF2000). The thawed PBMCs were resuspended in complete RPMI 1640, and additional cell viability and cell counts were performed on the resuspended cells. The PBMC cell suspension was divided into two primary aliquots. Cell viability and cell counts were measured for each primary aliquot. Primary aliquots were further subdivided into four sub-aliquots, each containing approximately 1 x 10^7^ cells, which were plated into individual wells of a 12-well cell culture plate. Two sub-aliquots served as duplicate controls and were treated with PBS, while the other two sub-aliquots were treated with 100 ng/mL of LPS (from *E. coli* 026:B6; eBioscience™ LPS, Catalog 00-4976, Invitrogen, Carlsbad, CA). All the cells were then incubated in a 37°C environment supplied with 5% CO for 24 hours. Following the 24-hour treatment period, cells were washed twice with warm RPMI 1640. Cell viability and cell counts were then measured. The cell suspensions were centrifuged at 1000g for 10 minutes, after which the cell supernatants and cell pellets were collected and stored at -80°C. The same technician performed all the experiments in one laboratory. All experiments were conducted in duplicate.

### 2.3 RNA isolation and sequencing

The PBMC pellets were retrieved from the -80°C freezer and thawed on a cool rack for approximately 15 minutes. Prior to RNA isolation, cell counts and cell viability were assessed for each sample on the Countess 3 FL. Each measurement was performed in duplicate, and the averages of the duplicate values were calculated to determine the total number of viable cells in each PBMC pellet. This information was used to assess the appropriate amount of lysis buffer required for RNA isolation. RNA was isolated from the PBMC pellets using the RNeasy mini kit (Qiagen, Catalog 74104) in accordance with the manufacturer’s instructions, which included a DNAse digestion step in the isolation protocol. Subsequently, the concentration, integrity, and purity of the isolated RNA were quantified using the ND 8000 spectrophotometer. The purified RNA was stored at -80°C. The RNA Clean & Concentrator-5 kit (Zymo Research, Catalog R1015) was later used for purification of the extracted RNA. The same technician performed all the RNA extractions and purifications in one laboratory. Total RNA from the three donors (12 total samples) was sent to the UC Davis Genomics Core Facility for library preparation and sequencing. Prior to library preparation, 200 nanograms (ng) of total RNA was treated with the QIAseq FastSelect–5S/16S/23S ribodepletion reagent (Qiagen, Hilden Germany) following the instructions of the manufacturer. (14–16) Dual-barcode indexed RNA-seq libraries were generated from 200 ng total RNA each using the Watchmaker RNA Library Prep Kit (Watchmaker Genomics, Boulder, Colorado). Eleven cycles of PCR amplification were used for double six base pair index addition and library fragment enrichment. The fragment size distribution of the libraries was verified via micro-capillary gel electrophoresis on TapeStation (Agilent, Santa Clara, CA). Prepared libraries were quantified on a Qubit fluorometer (LifeTechnologies, Carlsbad, CA) and pooled in equimolar ratios. Final pool was quantified using KAPA Illumina library quantitative PCR reagents (Roche Diagnostics Corp., Indianapolis, IN).

Sequencing of the 12 samples was done on an Illumina NovaSeqX (Illumina Inc., San Diego, CA). All 12 samples were multiplexed into a single pool, with each sample labeled with a dual-indexed adapter.(17) The sample pool was sequenced on four lanes for 150 cycles of paired-end reads with a 1% PhiX v3 control library spike (Illumina Inc., San Diego, CA). Post-sequencing basecall files (bclfiles) were demultiplexed and converted into a FASTQ file format using the bcl2fastq v2.17 software (Illumina Inc., San Diego, CA). Data were posted and retrieved from a secured FTP site hosted by the Core Facility.

### 2.4 RNA-seq Alignment, Data Processing, and Quality Control

RNA-seq data processing was performed based on best practices(18, 19) and our previous experience.(20, 21) Briefly, individual samples were inspected with FASTQC(22) and in aggregate with MultiQC.(23) Reads were filtered and trimmed using Trimmomatic.(24) The reference genome was prepared using the GRCh38.p14 assembly and transcriptome annotations were obtained from the Gencode v46 primary assembly.(25) Trimmed reads were aligned to the annotated reference genome using the STAR aligner.(26) Alignments were created for all RNA samples individually for gene level analysis (n=12) as well as combined across duplicates of a donor for alternative splicing analysis (n=6). Abundance of RNA at the gene level were estimated from the aligned reads as estimated counts using featureCounts(27) and as transcripts per million (TPM) using TPMcalculator.(28) Replicate count data from featureCounts were processed in edgeR and used for the gene level differential expression and pathway impact analyses.(29) Lower expressed tags were filtered out, and count estimates were normalized with the trimmed means of M-values (TMM) method.(30) Ensembl(31) gene and transcripts were annotated with Entrez gene ID and symbol.(32) Long non-coding RNAs were identified from the LNCipedia (version 5.2).(33) TPM estimates were used for cell-type composition deconvolution and alternative splicing analyses. Estimated count and TPM measures are available on NCBI GEO (Series GSE308443).

### 2.5 Global Differential Gene Expression Analysis

Differential expression was evaluated based on published best practices(34, 35) and our previous experience.(20, 36) Briefly, differential expression was determined under a variance modeling strategy that addressed the over-dispersion observed in gene expression count data using edgeR(37) (version 4.4.2).(38, 39) We assessed the significance of the transcriptome-wide analysis using a strict false discovery rate (FDR) of 5% under the Benjamini-Hochberg (BH) procedure.(40, 41) A heatmap of the top 50 differentially expressed genes was evaluated by clustering samples and genes using the aheatmap() function of the NMF R library (version 0.28).

We evaluated connectivity among differentially expressed genes and evaluated for functional enrichment of GO terms(42) using Search Tool for the Retrieval of Interacting Genes (STRING).(43) We assessed the significance of the protein-protein interaction (PPI) network enrichment using a p-value of <0.05. We also assessed for functional enrichment of differentially expressed genes in the GO terms using an FDR of 10% under the BH procedure. (40, 41)

### 2.6 Global Pathway Impact Analysis

Pathway analysis approaches traditionally consider pathways as lists of genes and ignore the additional information available in the pathway representation (e.g., topology). However, Pathway Impact Analysis (PIA) includes potentially important biological factors (e.g., gene-gene interactions, flow signals in a pathway, pathway topologies) as well as the magnitude (i.e., log fold-change) and the p-values from the DE analysis.(44) We used Pathway Express(45) to perform the PIA. The analysis included all genes (i.e., cutoff free) and the DE analysis results (i.e., p-value and log fold-change) to determine the probability of a pathway perturbation (pPERT). A total of 147 KEGG signaling pathways were evaluated.(46) Sequence loci data were annotated with Entrez gene IDs. The gene names were annotated using the HUGO Gene Nomenclature Committee resource database.(47) Significance of the pathway analyses was assessed using 10,000 permutations and a strict FDR of 5×10^-5^ under the BH procedure.(40, 41)

### 2.7 Cell Type Composition Estimates

Cell-type deconvolution was performed from TPM estimates from the quantTIseq(48) method using the immunedeconv R tool (49). Statistically significant differences between two groups were tested with two-sided Wilcoxon’s test. Significance was assessed using a nominal p-value of 0.05.

### 2.8 Differential Alternative Splicing Analyses

Analysis of differential regulation of RNA via splicing was performed using the alternative splicing mapping tool (rMATS-turbo)(50) following published protocols.(51) Loci were included in the analysis having an average read count ≥10 in both groups. Events with average percent spliced in (PSI) values <0.05 or >0.95 in both sample groups were included. We assessed for the significance of the differential splicing with a FDR cutoff of 0.01 and a between-group PSI value difference of |Δ PSI| ≥ 0.05. Heatmaps of the alternative splicing (AS) events were generated using Jutils (https://github.com/Splicebox/Jutils).(52)

### 2.9 Functional Characterization of Alternatively Spliced Genes

Functional enrichment for genes with differential alternative splicing was performed using the hypergeometric test for pathways in the BioPlanet, Elsevier, and WikiPathway databases for each event type. Significance of the enrichment was assessed with an FDR cutoff of 0.05 within each event.

Because an AS event may not be functional,(53, 54) or lead to different functions or interactions,(55) we used DIGGER (version 2.0)(56, 57) and Network-based Enrichment method for AS Events (NEASE)(58) to integrate pathways with structural annotations of protein-protein interactions to characterize the functional effects of AS events. Proteins with domains affected by changes in AS are first identified, followed by splicing-aware network enrichment analysis using an edge-level hypergeometric test to detect which pathways AS changes rewired interactions more often than by chance. All significant differentially alternatively spliced (DAS) skipped exon (SE) events are included in the analysis. Predicted protein domain-domain interactions (DDIs) with high-level confidence and divisible by 3 were included. Not-in-frame sites were excluded. We evaluated for enrichment of pathways from the Reactome and WikiPathways databases and assessed the significance of the enrichment with an FDR cutoff of 0.05.

## 3 Results

### 3.1 Global Differential Expression and Pathway Perturbation between LPS and PBS-treated PBMCs

Samples ranged between 59.0 - 76.5 M uniquely aligned reads. When we looked at the gene expression patterns in an unsupervised manner using PCA, we saw differences between the LPS and PBS treated groups (Supplemental Figure 1). Of the 15,879 genes evaluated, 490 were significantly differentially expressed (Table 1, Supplemental File 2, Supplemental Figure 2). Table 1 presents the top 20 differentially expressed genes in human PBMCs between LPS stimulated samples and PBS controls. A heatmap of expression of the top 50 differentially expressed genes is shown in Figure 1.

**Table 1.**
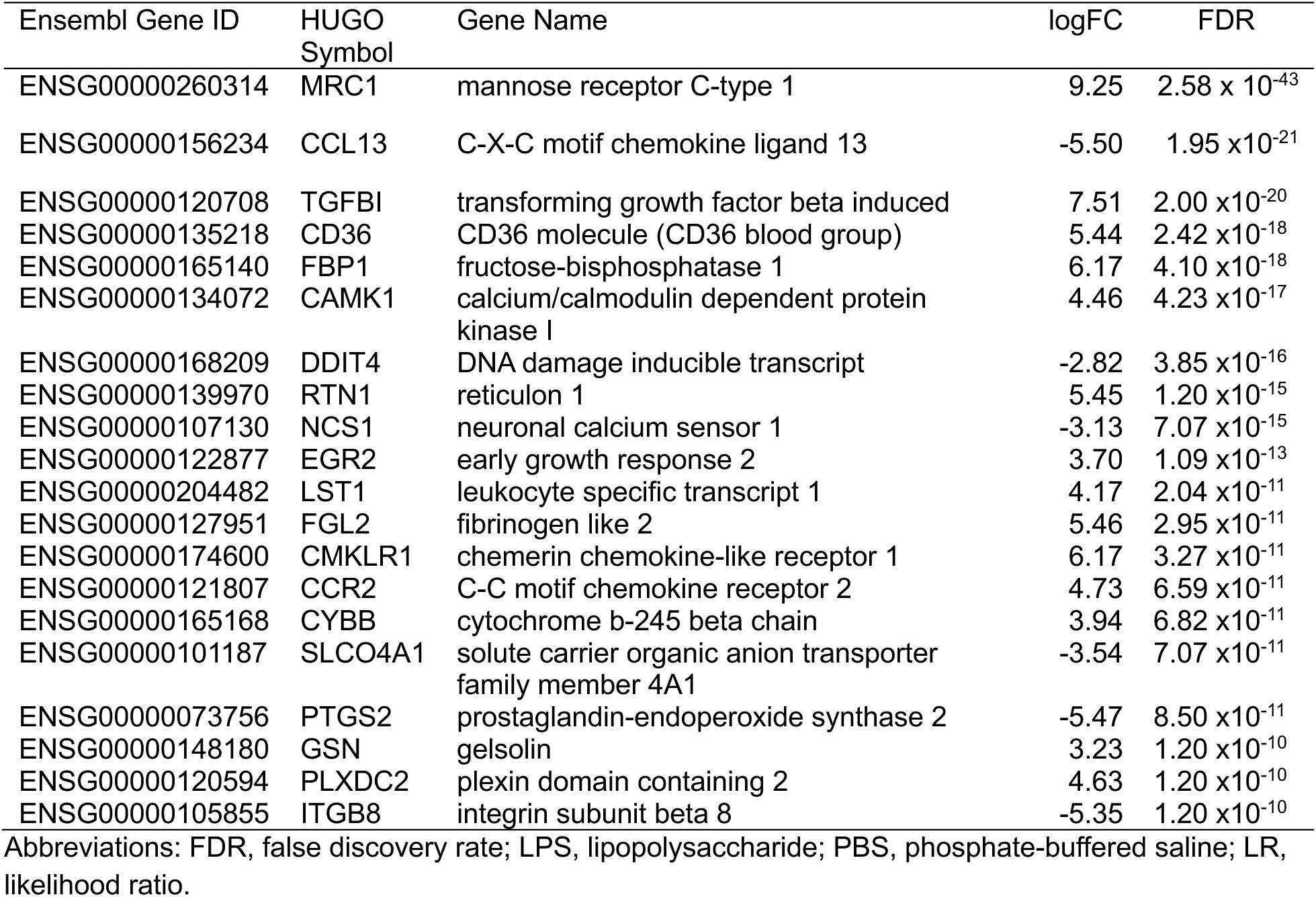
Top 20 differentially expressed genes in human PBMCs between LPS stimulated and PBS controls.

**Figure 1.**
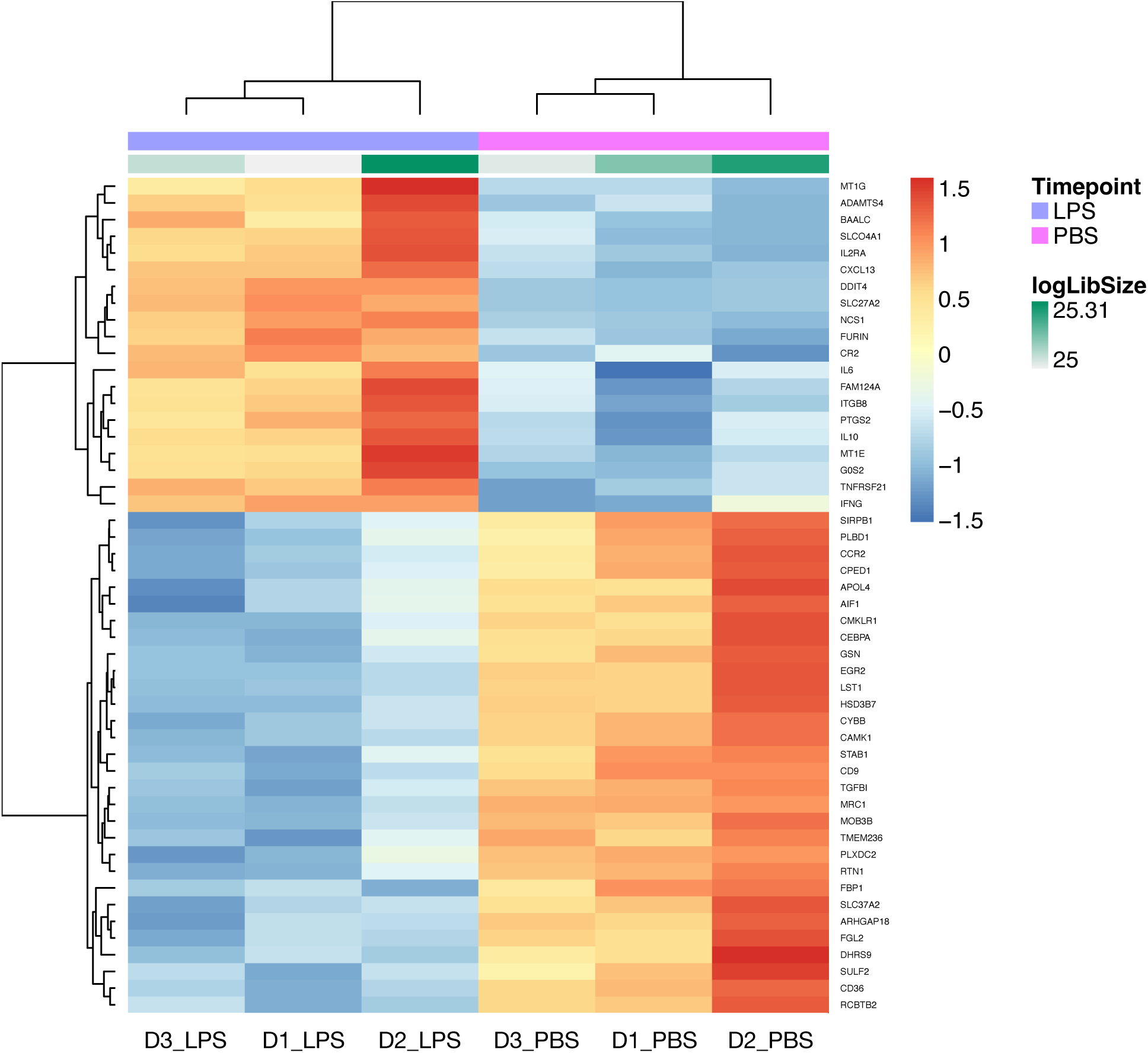
Heatmap of the top 50 differentially expressed genes between LPS and PBS-treated PBMCs from three women donors identified in the global transcriptome analysis. Rows and columns were clustered using the Euclidean distance and the complete linkage method.

We identified 446 differentially expressed genes annotated for the HUGO symbol to be included in the functional analysis with STRING. The resulting network of 446 genes (Figure 2) had significantly more interactions than expected for a random set of proteins of similar size drawn from the genome (number of edges=878, average node degree=3.94, avg. clustering coefficient=0.362, expected number of edges=215, PPI enrichment p-value <1.0×10^-16^). Significant enrichment was found for 646 GO terms (n=560 biological processes, n=58 cellular component, n=28 molecular function)(Supplemental Figure 3, Supplemental File 3).

**Figure 2.**
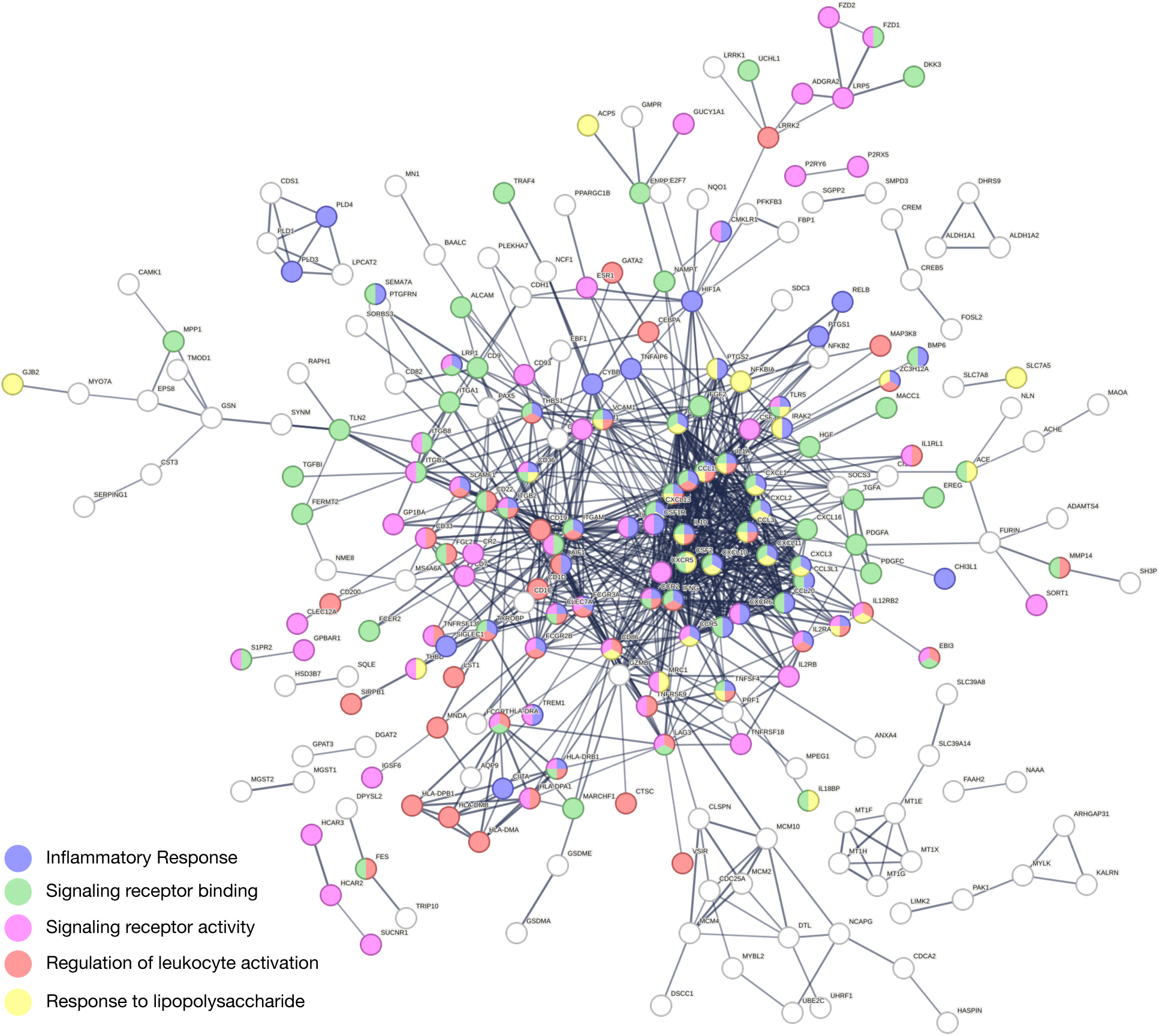
STRING connectivity network demonstrating a protein-protein interaction network of predicted functional partners for differentially expressed genes. Nodes represent all proteins produced by a single protein-coding gene locus. Disconnected nodes in the network are not displayed.

Of the 147 KEGG pathways evaluated, 46 were perturbed (Table 2, Supplemental File 4). 1Perturbed signaling pathways were found for biological functions related to fundamental cellular signaling and ion dynamics, cellular growth, development, and survival, immune system regulation and host defense, hormonal and systemic physiological processes, and metabolic control and energy homeostasis (Table 2). These findings demonstrate our study accurately recapitulates the known global transcriptional response to LPS stimulation.

**Table 2.** Perturbed KEGG signaling pathways associated with LPS as compared to PBS treated human PBMCs in the global transcriptome analysis.

| Pathway ID | Pathway name | Pert. Score | pPert |
| --- | --- | --- | --- |
| **Fundamental Cellular Signaling and Ion Dynamics** |  |  |  |
| hsa03013 | RNA transport | 33.45 | 0.00003 |
| hsa04144 | Endocytosis | 22.69 | 0.00003 |
| hsa04540 | Gap junction | 20.05 | 0.00003 |
| hsa04122 | Sulfur relay system | 18.42 | 0.00003 |
| hsa03015 | mRNA surveillance pathway | 17.59 | 0.00003 |
| hsa03008 | Ribosome biogenesis in eukaryotes | 17.01 | 0.00003 |
| hsa03018 | RNA degradation | 16.02 | 0.00003 |
| hsa04020 | Calcium signaling pathway | 15.39 | 0.00003 |
| hsa04141 | Protein processing in endoplasmic reticulum | 13.91 | 0.00003 |
| hsa04130 | SNARE interactions in vesicular transport | 11.64 | 0.00003 |
| **Cellular Growth, Development, and Survival** |  |  |  |
| hsa04520 | Adherens junction | 26.13 | 0.00003 |
| hsa04010 | MAPK signaling pathway | 25.32 | 0.00003 |
| hsa04320 | Dorso-ventral axis formation | 22.54 | 0.00003 |
| hsa04350 | TGF-beta signaling pathway | 21.48 | 0.00003 |
| hsa04151 | PI3K-Akt signaling pathway | 19.09 | 0.00003 |
| hsa04370 | VEGF signaling pathway | 18.04 | 0.00003 |
| hsa04510 | Focal adhesion | 17.80 | 0.00003 |
| hsa04340 | Hedgehog signaling pathway | 17.13 | 0.00003 |
| hsa04514 | Cell adhesion molecules (CAMs) | 16.99 | 0.00003 |
| hsa04530 | Tight junction | 16.25 | 0.00003 |
| hsa04210 | Apoptosis | 16.18 | 0.00003 |
| hsa04360 | Axon guidance | 15.48 | 0.00003 |
| hsa04012 | ErbB signaling pathway | 15.19 | 0.00003 |
| hsa04390 | Hippo signaling pathway | 14.37 | 0.00003 |
| hsa04115 | p53 signaling pathway | 14.29 | 0.00003 |
| hsa04330 | Notch signaling pathway | 14.15 | 0.00003 |
| hsa04612 | Antigen processing and presentation | 13.78 | 0.00003 |
| hsa03460 | Fanconi anemia pathway | 13.55 | 0.00003 |
| hsa04512 | ECM-receptor interaction | 13.21 | 0.00003 |
| hsa04110 | Cell cycle | 13.18 | 0.00003 |
| hsa04310 | Wnt signaling pathway | 12.58 | 0.00003 |
| **Immune System Regulation and Host Defense** |  |  |  |
| hsa04145 | Phagosome | 20.14 | 0.00003 |
| hsa04610 | Complement and coagulation cascades | 16.21 | 0.00003 |
| hsa04064 | NF-kappa B signaling pathway | 15.95 | 0.00003 |
| hsa04062 | Chemokine signaling pathway | 15.66 | 0.00003 |
| hsa04060 | Cytokine-cytokine receptor interaction | 14.10 | 0.00003 |
| hsa04620 | Toll-like receptor signaling pathway | 13.99 | 0.00003 |
| **Hormonal and Systemic Physiological Processes** |  |  |  |
| hsa04080 | Neuroactive ligand-receptor interaction | 19.48 | 0.00003 |
| hsa04270 | Vascular smooth muscle contraction | 22.50 | 0.00003 |
| hsa04260 | Cardiac muscle contraction | 18.02 | 0.00003 |
| hsa04114 | Oocyte meiosis | 13.55 | 0.00003 |
| hsa04380 | Osteoclast differentiation | 13.16 | 0.00003 |
| **Metabolic Control and Energy Homeostasis** |  |  |  |
| hsa04140 | Regulation of autophagy | 18.98 | 0.00003 |
| hsa04150 | mTOR signaling pathway | 18.70 | 0.00003 |
| hsa03320 | PPAR signaling pathway | 15.21 | 0.00003 |
| hsa04066 | HIF-1 signaling pathway | 14.72 | 0.00003 |

Abbreviations: LPS, lipopolysaccharide; PBS, phosphate-buffered saline; Pert, normalized perturbation score; pPert, false discovery rate adjusted perturbation p-value.

### 3.2 Cell type compositions associated with LPS treatment

Cell type estimates were obtained for ten cell types (Table 3). LPS stimulation of PBMCs is associated with increased levels of Macrophage M1 and reduced levels of Macrophage M2 (Figure 3A). No differences were found in Natural Killer, Myeloid Dendritic, Monocyte, Neutrophil, B or any T cell types.

**Figure 3.**
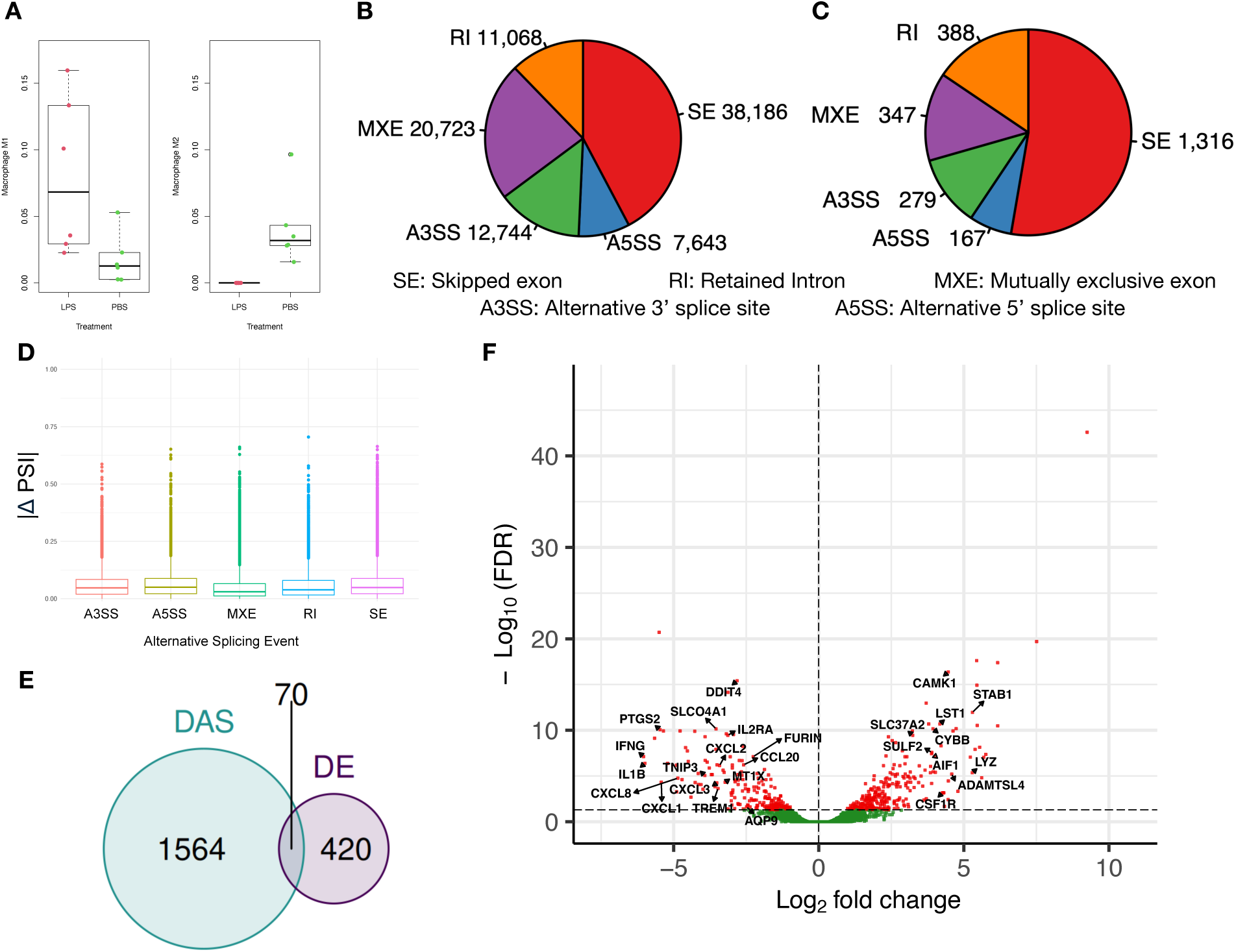
(A) Differences in the abundance of Macrophage M1 and M2 cell types by treatment estimated by immune deconvolution (quanTIseq). (B) Summary pie chart of total alternative splicing events identified in the LPS- and PBS-treated cells after filtering for high read support and extreme PSI values. (C) Summary pie chart of differential alternative splicing events between the LPS and PBS treatments (FDR <0.01, |deltaPsi| ≥0.05). (D) Box plots of the absolute value of deltaPSI by event type. (E) A Venn diagram depicts the overlap of differentially expressed (DE) genes and differentially alternatively spliced (DAS) genes across all splicing events. (F) Volcano plot of DE genes between LPS and PBS-treated donor PBMC samples. Significantly DE genes are colored red (FDR < 0.05), non-significant genes are colored green. Space permitting, a number of genes that are both DE and DAS are labeled for reference.

**Table 3.**
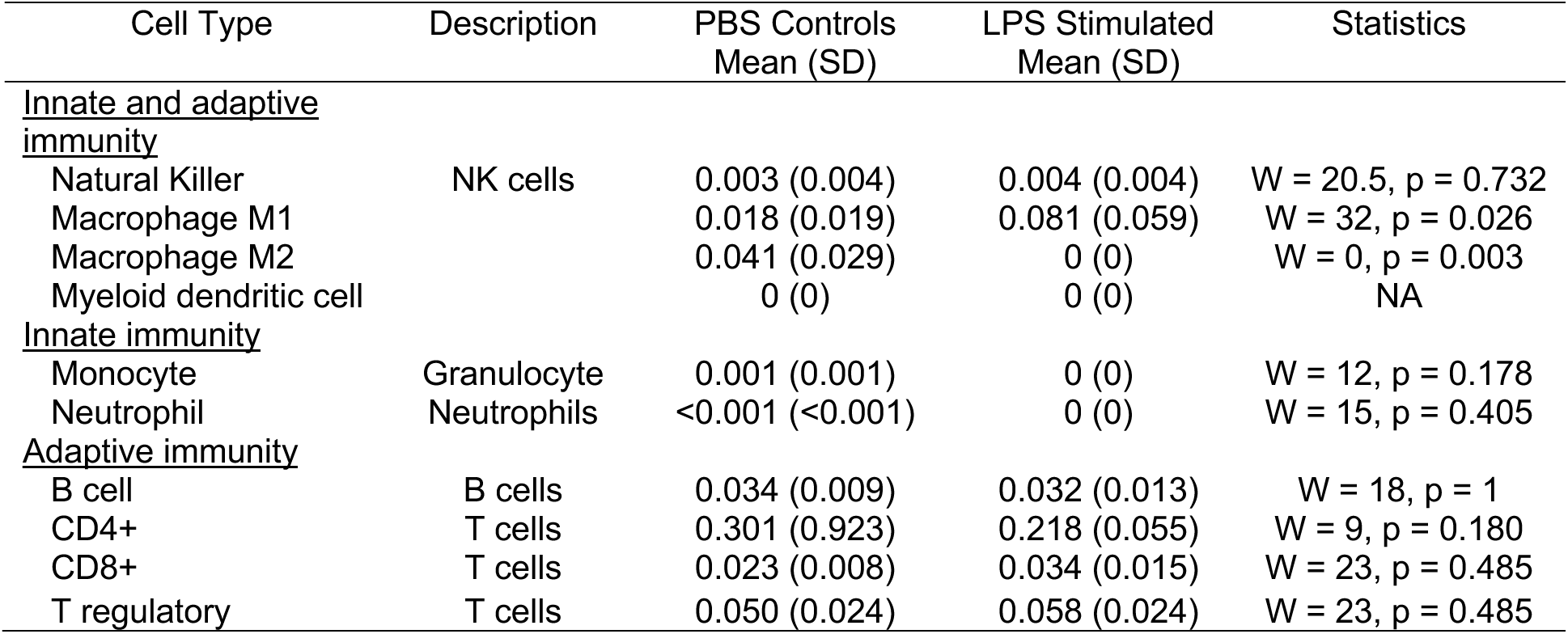
Differences in Transcriptomic Estimates of Cell Type Compositions Proportions Between LPS and PBS Treated PBMCs

| Cell Type | Description | PBS Controls<br>Mean (SD) | LPS Stimulated<br>Mean (SD) | Statistics |
| --- | --- | --- | --- | --- |
| <u>Innate and adaptive immunity</u> |  |  |  |  |
| Natural Killer | NK cells | 0.003 (0.004) | 0.004 (0.004) | W = 20.5, p = 0.732 |
| Macrophage M1 |  | 0.018 (0.019) | 0.081 (0.059) | W = 32, p = 0.026 |
| Macrophage M2 |  | 0.041 (0.029) | 0 (0) | W = 0, p = 0.003 |
| Myeloid dendritic cell |  | 0 (0) | 0 (0) | NA |
| <u>Innate immunity</u> |  |  |  |  |
| Monocyte | Granulocyte | 0.001 (0.001) | 0 (0) | W = 12, p = 0.178 |
| Neutrophil | Neutrophils | <0.001 (<0.001) | 0 (0) | W = 15, p = 0.405 |
| <u>Adaptive immunity</u> |  |  |  |  |
| B cell | B cells | 0.034 (0.009) | 0.032 (0.013) | W = 18, p = 1 |
| CD4+ | T cells | 0.301 (0.923) | 0.218 (0.055) | W = 9, p = 0.180 |
| CD8+ | T cells | 0.023 (0.008) | 0.034 (0.015) | W = 23, p = 0.485 |
| T regulatory | T cells | 0.050 (0.024) | 0.058 (0.024) | W = 23, p = 0.485 |

### 3.3 Differential Alternative Splicing and Pathway Enrichment is Associated with LPS treatment in PBMCs

Sample read coverage ranged between 128.7M to 143.3M uniquely aligned reads, a range that has been recently identified to be sufficient to adequately identify highly expressed genes.(59) A total of 338,041 splice events were evaluated (Supplemental Table 2). After filtering, 90,364 splice events were identified in the LPS and PBS-treated PBMCs (Figure 3B, Supplemental File 5). Of these, 1,316 skipped exon (SE), 388 retained intron (RI), 347 mutually exclusive exon (MXE), 279 alternative 3’ splice (A3SS), and 167 alternative 5’ splice (A5SS) site events were significant (Figure 3C). A total of 1,634 unique genes were identified across the 2,507 splice events. The SE events showed a similar range of Δ PSI as compared to the other events (Figure 3D). Of these, 70 genes were both differentially expressed and differentially alternatively spliced (Figures 3E, 3F). A heatmap of the top 50 differentially alternatively spliced events is shown in Figure 4.

**Figure 4.**
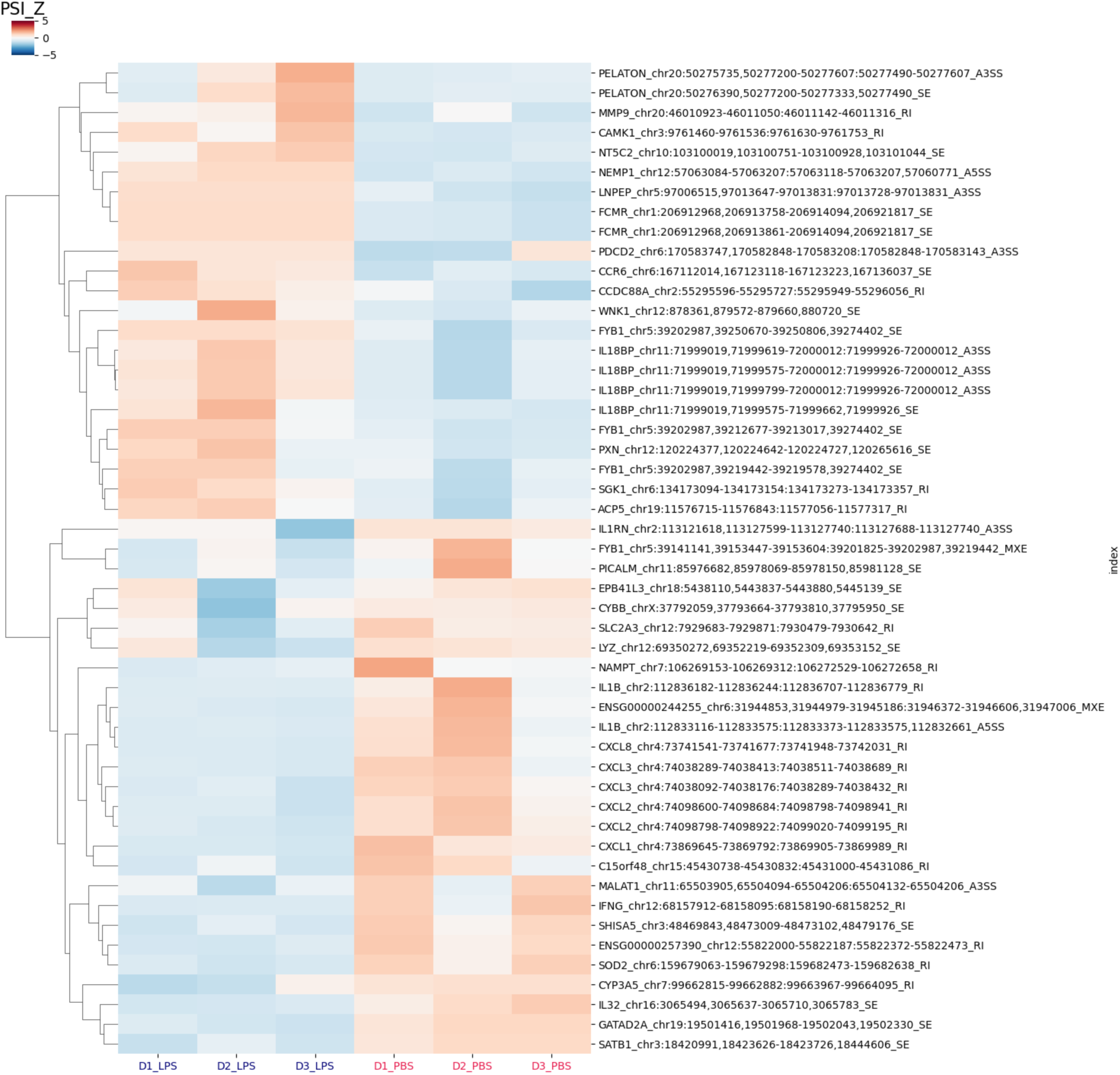
Heatmap of top differentially alternatively spliced events (FDR < 10^-6^) between lipopolysaccharide (LPS) or phosphate-buffered saline (PBS) treated PBMCs from three women donors (D). Supervised weighted clustering was performed with the cityblock metric.

The summary of the enrichment tests for each pathway database for each event type is reported in Supplemental Table 3. The volcano plots and top 10 enriched pathways from each database for each event are shown in Figure 5. Of the 1,316 SE events, 698 were downregulated and 618 were upregulated in the LPS-treated PBMCs. In terms of SE events, of the 1071 BioPlanet pathways evaluated, 139 were enriched. Of the 1190 Elsevier pathways evaluated, 132 were enriched. Of the 446 WikiPathway pathways evaluated, 58 were enriched. The results of the enrichment tests for all site event types are provided in Supplemental File 6.

**Figure 5.**
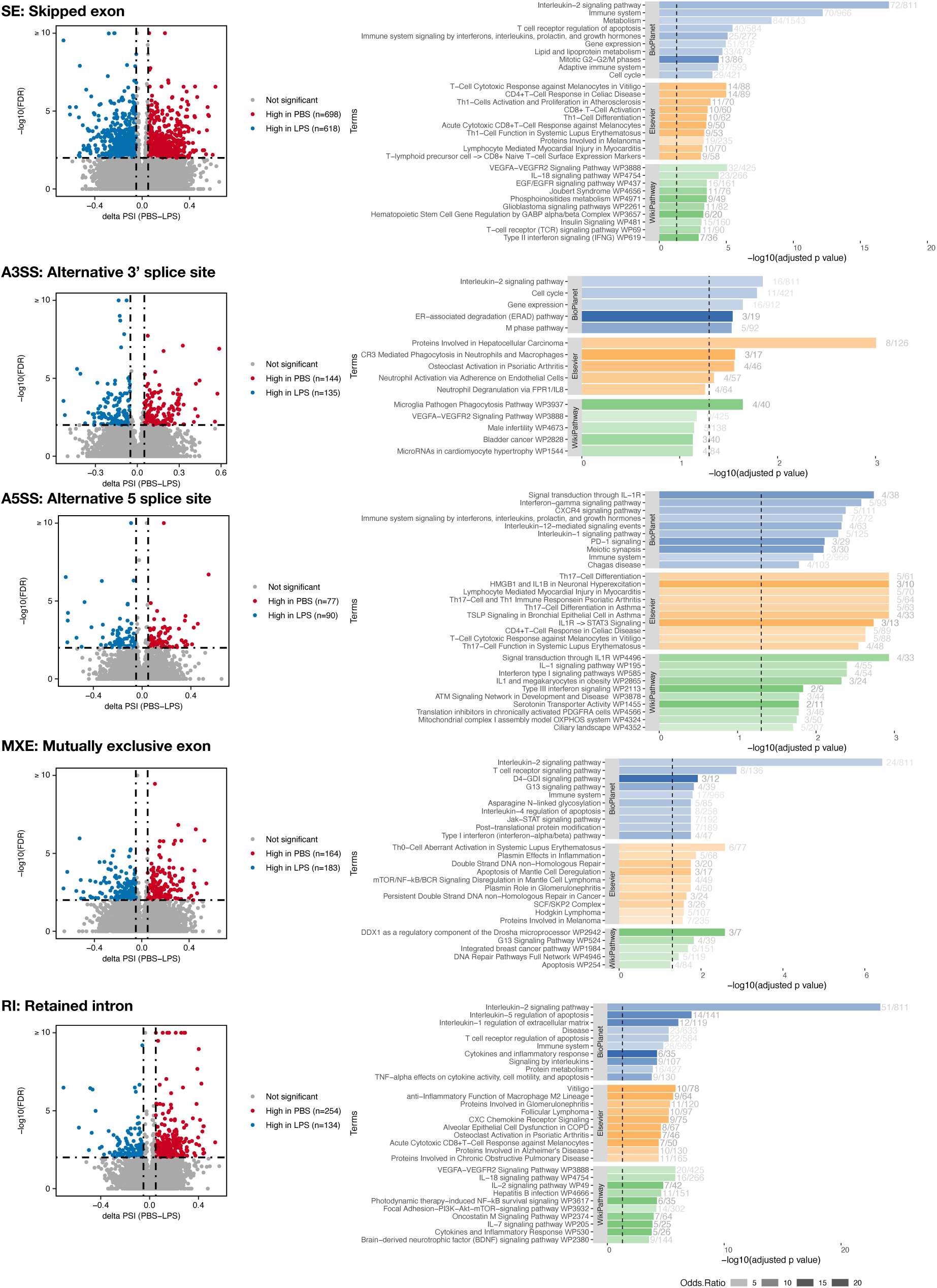
Volcano and pathway enrichment plots for each splicing event type. The bar graphs show the top ten enriched ways for the BioPlanet, Elsevier, and WikiPathway databases. The bar lengths depict the Benjamini-Hochberg adjusted P values calculated from a hypergeometric test. The odds ratio of enrichment is indicated by the opacity of bars. numbers shown beside each bar represent the number of differentially spliced genes and the total number of genes in the corresponding pathway, respectively.

A total of 156 DAS events (A3SS n=10, A5SS n=9, RI n=19, MXE n=28, SE n=90) were identified across 88 distinct long non-coding RNA (lncRNA) genes (Table 4). Of these genes, multiple DAS events were identified for 30 genes, and one gene (ENSG00000293154) also had global differential expression.

**Table 4.**
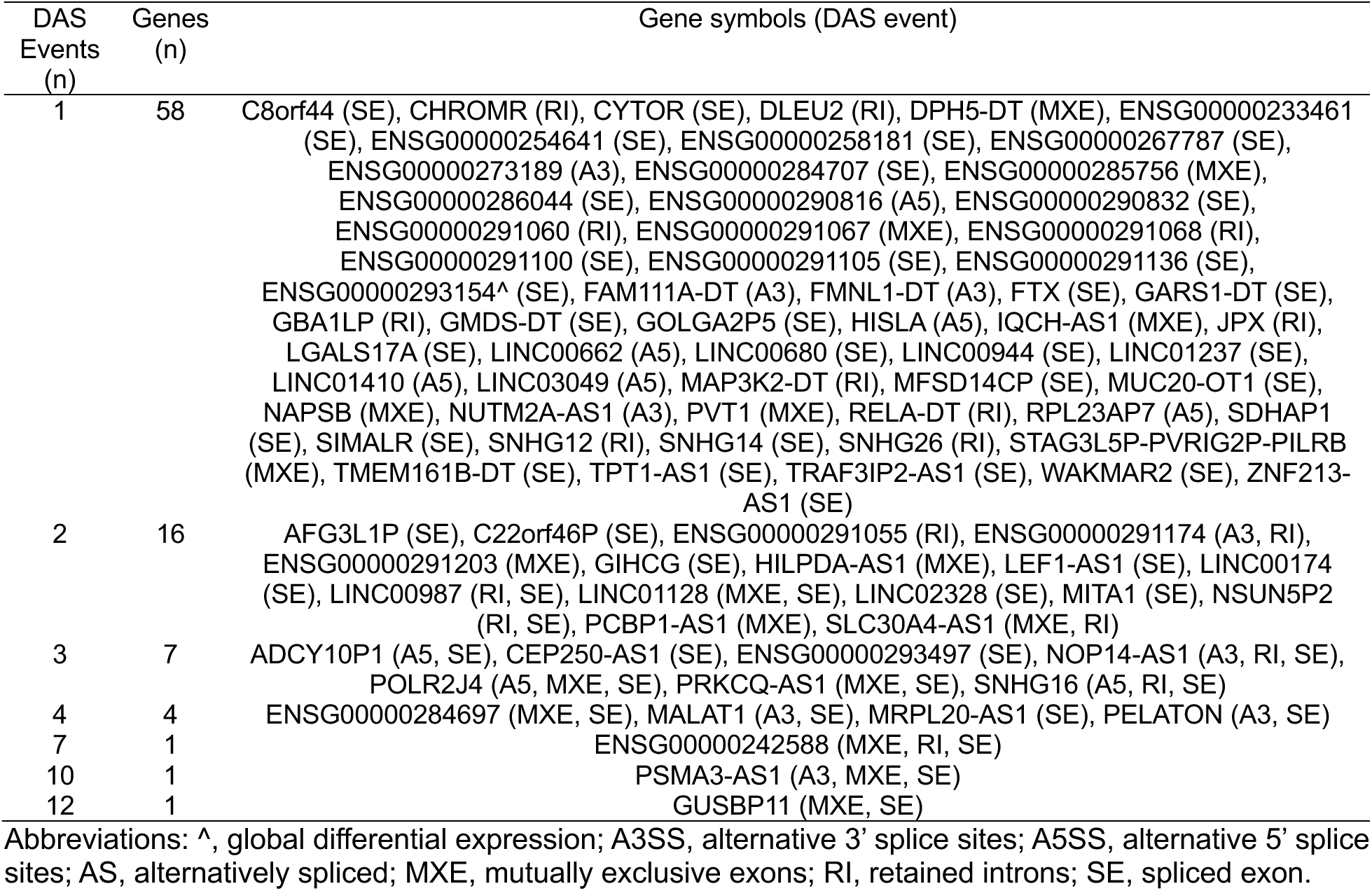
A summary of the annotated lncRNA genes that were identified to have differential alternative splicing in human PBMCs stimulated with LPS as compared to PBS.

### 3.4 Splice Aware Network Interaction Functional Enrichment of Alternative Splicing Events Associated with LPS Treatment in PBMCs

A total of 1,316 SE events were included in the analysis. The analysis identified 138 affected protein domains, 2 affected linear motifs, 24 affected interacting (with residue evidence), 72 affected domains/motifs with known interactions, and 1,596 affected protein interactions/bindings. An effect of alternative splicing on protein features was identified for 14% of AS genes (Supplemental Figure 4A). Of these, the majority of the affected features were protein domains (83%), followed by residues (15.0%) and linear motifs (1.0%) (Supplemental Figure 4B) Splice-aware network analysis identified 70 Reactome and 48 WikiPathway pathways significantly enriched for AS exons (Figure 6, Supplemental File 7). Enriched pathways were found for six categories of biological functions related to Immune system and inflammation (e.g., Regulation of toll-like receptor signaling pathway, Table 5), Signaling pathways (E.g., Negative regulation of the PI3K/AKT network, Table 6), Cell adhesion, cytoskeleton and extracellular matrix (e.g., Focal adhesion), Metabolism and biosynthesis (e.g., Glycosaminoglycan metabolism), Cellular processes and vesicle trafficking (e.g., Neddylation), and Cancer and disease pathways (e.g., Diseases of signal transduction, Table 7). A summary of the alternatively spliced genes (e.g., TLR4, MAP2K2) and their predicted targets (e.g., MYD88, IRAK2) for the Negative regulation of the PI3K/AKT network pathway are reported in Table 8.

**Figure 6.**
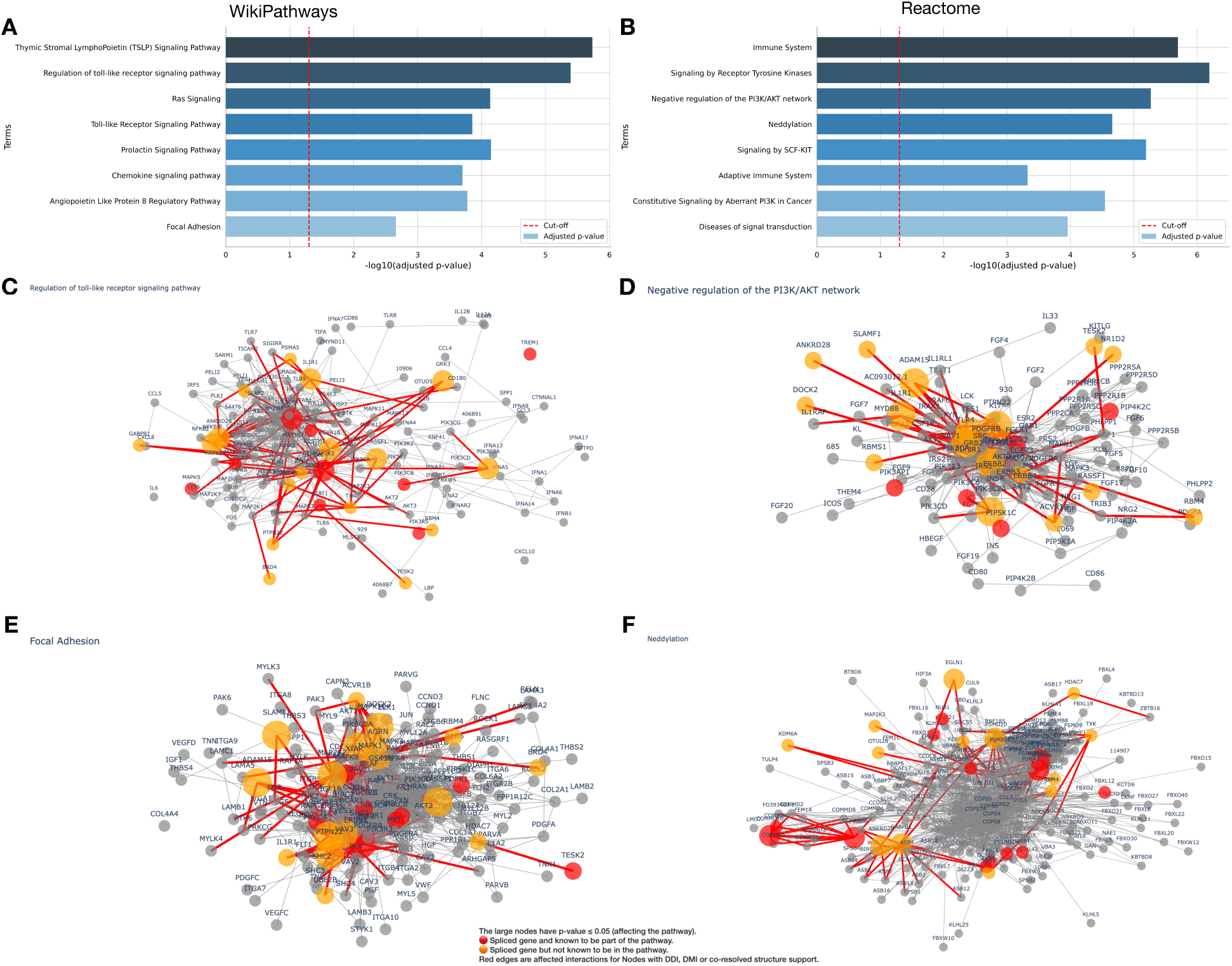
Splice-aware pathway protein-protein interaction (PPI) network enrichment analysis of exon skipping events. Top eight enriched pathways from the (A) WikiPathway and (B) Reactome databases. Visualization of the spliced genes on the PPI network for the pathways: (C) Regulation of toll-like receptor signaling pathway (D) Negative regulation of the PI3K/AKT network, (E) Focal Adhesion, and (F) Neddylation.

**Table 5.** Immune System and Inflammation pathways enriched for differentially alternatively spliced genes of human PBMCs ted with LPS or PBS predicted to impact protein-protein network interactions.

| Pathway ID | Pathway name | FDR | NEASE score |
| --- | --- | --- | --- |
| WP1449 | Regulation of toll-like receptor signaling pathway | 4.04E-06 | 23.74 |
| WP75 | Toll-like Receptor Signaling Pathway | 7.31E-05 | 17.69 |
| R-HSA-202403 | TCR signaling | 1.55E-04 | 16.80 |
| WP3929 | Chemokine signaling pathway | 1.39E-04 | 15.89 |
| R-HSA-512988 | Interleukin-3, 5 and GM-CSF signaling | 2.87E-04 | 15.87 |
| R-HSA-1280218 | Adaptive Immune System | 4.74E-04 | 18.58 |
| WP4197 | The human immune response to tuberculosis | 7.26E-04 | 13.24 |
| WP585 | Interferon type I signaling pathways | 1.08E-03 | 12.98 |
| R-HSA-6785807 | Interleukin-4 and 13 signaling | 1.85E-03 | 9.29 |
| WP395 | IL-4 Signaling Pathway | 2.70E-03 | 10.12 |
| WP195 | IL-1 signaling pathway | 3.61E-03 | 9.76 |
| WP3877 | Simplified Depiction of MYD88 Distinct Input-Output Pathway | 4.38E-03 | 9.38 |
| WP286 | IL-3 Signaling Pathway | 4.81E-03 | 9.05 |
| R-HSA-168249 | Innate Immune System | 9.76E-03 | 12.58 |
| WP2865 | IL1 and megakaryocytes in obesity | 6.56E-03 | 8.67 |
| R-HSA-877312 | Regulation of IFNG signaling | 1.47E-02 | 6.19 |
| R-HSA-389356 | CD28 co-stimulation | 1.70E-02 | 8.47 |
| R-HSA-202430 | Translocation of ZAP-70 to Immunological synapse | 1.77E-02 | 6.83 |
| R-HSA-2871809 | FCERI mediated Ca+2 mobilization | 1.88E-02 | 5.81 |
| WP23 | B Cell Receptor Signaling Pathway | 1.25E-02 | 8.34 |
| R-HSA-388841 | Costimulation by the CD28 family | 2.32E-02 | 7.28 |
| R-HSA-1280215 | Cytokine Signaling in Immune system | 2.69E-02 | 9.41 |
| R-HSA-1810476 | RIP-mediated NFkB activation via ZBP1 | 2.69E-02 | 7.01 |
| R-HSA-202427 | Phosphorylation of CD3 and TCR zeta chains | 2.75E-02 | 6.23 |
| R-HSA-445989 | TAK1 activates NFkB by phosphorylation and activation of IKKs complex | 3.03E-02 | 6.85 |
| R-HSA-451927 | Interleukin-2 family signaling | 3.06E-02 | 7.47 |
| R-HSA-975110 | TRAF6 mediated IRF7 activation in TLR7/8 or 9 signaling | 3.52E-02 | 5.95 |
| R-HSA-912694 | Regulation of IFNA signaling | 3.59E-02 | 5.11 |
| WP49 | IL-2 Signaling Pathway | 2.65E-02 | 6.56 |
| WP3937 | Microglia Pathogen Phagocytosis Pathway | 2.94E-02 | 6.38 |
| R-HSA-2454202 | Fc epsilon receptor (FCERI) signaling | 3.99E-02 | 6.96 |
| WP69 | T-Cell antigen Receptor (TCR) Signaling Pathway | 3.33E-02 | 6.08 |
| WP619 | Type II interferon signaling (IFNG) | 4.70E-02 | 5.20 |
| WP4478 | LTF danger signal response pathway | 4.92E-02 | 4.66 |

Abbreviations: AS, alternative splicing; FDR, false discovery rate; LPS, lipopolysaccharide; NEASE, Network-based Enrichment method for AS Events; PBS, phosphate-buffered saline.

**Table 6.** Signaling pathways enriched for differentially alternatively spliced genes of human PBMCs treated with LPS or PBS predicted to impact protein-protein network interactions.

| Pathway ID | Pathway name | FDR | NEASE score |
| --- | --- | --- | --- |
| R-HSA-9006934 | Signaling by Receptor Tyrosine Kinases | 6.41E-07 | 29.84 |
| R-HSA-982772 | Growth hormone receptor signaling | 1.02E-06 | 17.87 |
| WP2203 | Thymic Stromal Lymphopoietin (TSLP) Signaling Pathway | 1.84E-06 | 24.20 |
| R-HSA-199418 | Negative regulation of the PI3K/AKT network | 5.40E-06 | 20.93 |
| R-HSA-1433557 | Signaling by SCF-KIT | 6.39E-06 | 18.96 |
| R-HSA-2219530 | Constitutive Signaling by Aberrant PI3K in Cancer | 2.84E-05 | 18.38 |
| WP4223 | Ras Signaling | 7.09E-05 | 17.97 |
| R-HSA-6811558 | PI5P, PP2A and IER3 Regulate PI3K/AKT Signaling | 8.57E-05 | 16.82 |
| R-HSA-195399 | VEGF binds to VEGFR leading to receptor dimerization | 9.63E-05 | 10.77 |
| R-HSA-194313 | VEGF ligand-receptor interactions | 9.63E-05 | 10.77 |
| WP2037 | Prolactin Signaling Pathway | 1.39E-04 | 17.18 |
| R-HSA-2586552 | Signaling by Leptin | 1.91E-04 | 11.63 |
| WP3915 | Angiopoietin Like Protein 8 Regulatory Pathway | 1.67E-04 | 15.22 |
| WP581 | EPO Receptor Signaling | 3.33E-04 | 13.24 |
| R-HSA-194306 | Neurophilin interactions with VEGF and VEGFR | 1.74E-03 | 8.14 |
| WP304 | Kit receptor signaling pathway | 1.08E-03 | 13.12 |
| WP3972 | PDGFR-beta pathway | 1.71E-03 | 10.80 |
| WP437 | EGF-EGFR Signaling Pathway | 2.19E-03 | 10.30 |
| R-HSA-202433 | Generation of second messenger molecules | 9.26E-03 | 8.60 |
| R-HSA-109704 | PI3K Cascade | 1.13E-02 | 9.06 |
| WP2593 | JAK-STAT | 6.12E-03 | 8.75 |
| R-HSA-1433559 | Regulation of KIT signaling | 1.50E-02 | 8.69 |
| R-HSA-8874081 | MET activates PTK2 signaling | 1.57E-02 | 7.86 |
| R-HSA-5673001 | RAF/MAP kinase cascade | 1.72E-02 | 8.43 |
| R-HSA-449147 | Signaling by Interleukins | 1.88E-02 | 11.13 |
| R-HSA-5684996 | MAPK1/MAPK3 signaling | 2.69E-02 | 7.72 |
| WP3932 | Focal Adhesion-PI3K-Akt-mTOR-signaling pathway | 2.27E-02 | 6.58 |
| WP2380 | Brain-Derived Neurotrophic Factor (BDNF) signaling pathway | 4.73E-02 | 5.18 |
| R-HSA-194138 | Signaling by VEGF | 3.52E-02 | 8.92 |
| R-HSA-8875513 | MET interacts with TNS proteins | 3.59E-02 | 5.90 |
| R-HSA-6802957 | Oncogenic MAPK signaling | 3.59E-02 | 5.87 |
| WP4172 | PI3K-Akt Signaling Pathway | 2.73E-02 | 6.54 |
| R-HSA-6802952 | Signaling by BRAF and RAF fusions | 3.95E-02 | 4.94 |
| R-HSA-186763 | Downstream signal transduction | 4.27E-02 | 7.39 |
| R-HSA-392517 | Rap1 signalling | 4.46E-02 | 5.53 |
| WP382 | MAPK Signaling Pathway | 4.59E-02 | 5.22 |
| WP2380 | Brain-Derived Neurotrophic Factor (BDNF) signaling pathway | 4.73E-02 | 5.18 |
| WP673 | ErbB Signaling Pathway | 4.81E-02 | 5.08 |
| WP3888 | VEGFA-VEGFR2 Signaling Pathway | 4.85E-02 | 4.99 |

Abbreviations: AS, alternative splicing; FDR, false discovery rate; LPS, lipopolysaccharide; NEASE, Network-based Enrichment method for AS Events; PBS, phosphate-buffered saline.

**Table 7.**
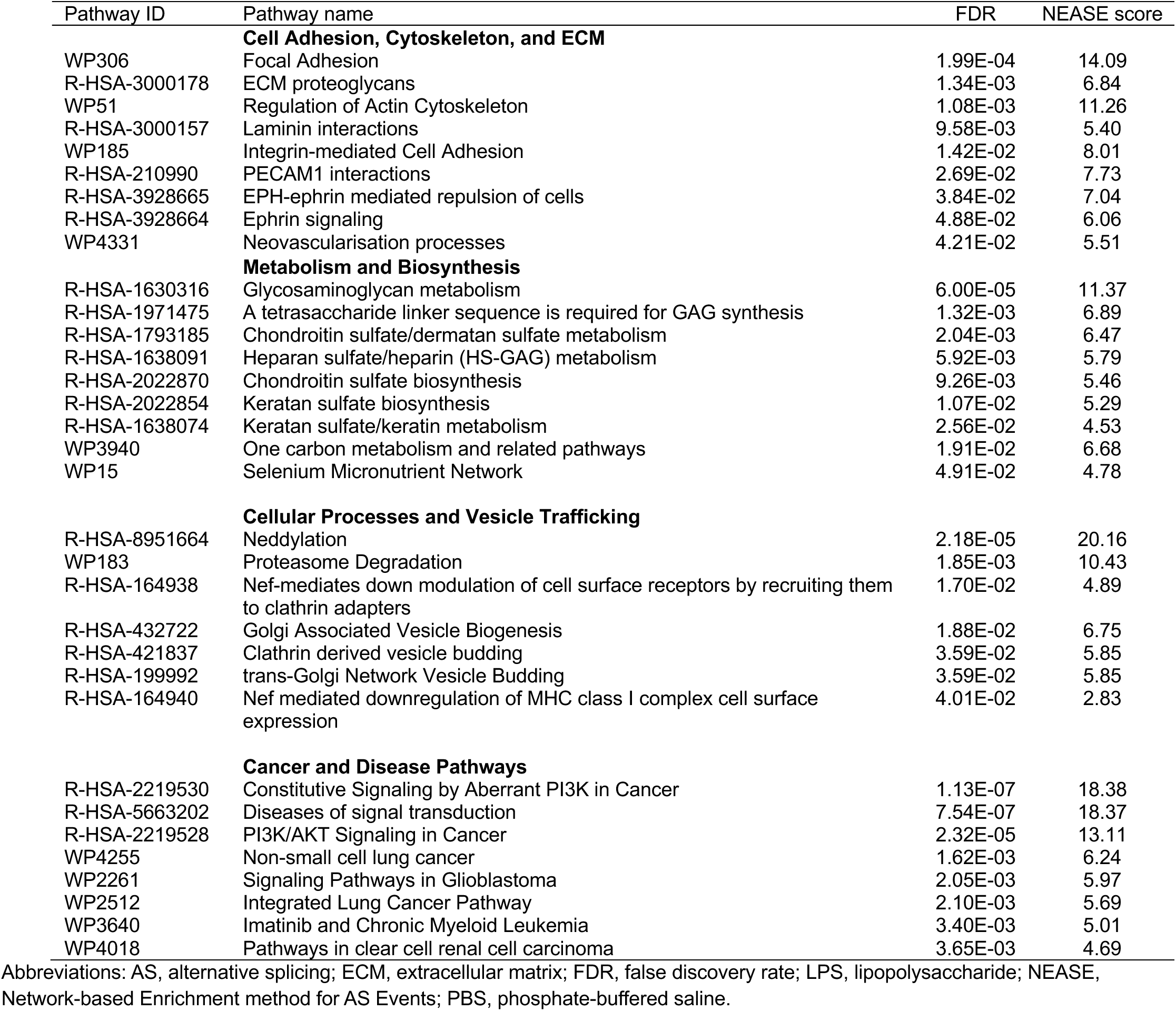
Pathways enriched for differentially alternatively spliced genes of human PBMCs treated with LPS or PBS predicted to protein-protein network interactions not primarily categorized as either immune response or signaling.

**Table 8.** Summary of alternatively spliced genes between LPS and PBS-treated PBMCs, driving the enrichment of the Regulation of Toll-like Receptor Signaling pathway (WP1449) and the genes affected by alternative splicing in the NEASE analysis.

| Spliced gene | Number of edges associated with the pathway (%) | P-value | Affected binding (edges) |
| --- | --- | --- | --- |
| TLR4* | 16/95 (16.84%) | 4.16x10<sup>-08</sup> | AC093012.1, TLR7, CD180, MYD88, TLR5, MAP3K7, SIGIRR, SYK, TLR9, RIPK1, TLR2, BTK, TLR1, TLR3, TLR4, LY96 |
| NFKBIE | 9/33 (27.27%) | 5.41x10<sup>-07</sup> | IRAK3, RIPK1, NFKBIA, MYD88, IRAK2, RELA, NFKB1, NFKB2, IRAK1 |
| ANKRD28 | 8/65 (12.31%) | 9.23x10<sup>-04</sup> | PLK1, IRAK3, RIPK1, NFKBIA, TBK1, IRAK2, RELA, IRAK1 |
| ACVR1B | 8/61 (13.11%) | 6.01x10<sup>-04</sup> | SMAD6, MAPK13, MAPK10, SQSTM1, MAPK3, MAPK12, MAPK1, MAPK11 |
| IL1R1 | 4/9 (44.44%) | 1.05x10<sup>-04</sup> | PIK3R1, SIGIRR, MYD88, IL1B |
| MAP2K2* | 17/199 (8.54%) | 1.89x10<sup>-04</sup> | MAPK10, IRAK3, MAPK8, IRAK2, AKT3, MAP2K3, MAPK13, MAPK1, MAP2K6, MAPK3, AKT2, MAPK11, MAP3K8, MAPK12, MAP2K2, IRAK1, MAP2K1 |
| PIK3C2A | 4/11 (36.36%) | 2.61x10<sup>-04</sup> | PIK3CB, PIK3CD, PIK3CA, PIK3CG |
| GRK3 | 3/10 (30.00%) | 3.09x10<sup>-03</sup> | MAPK13, MAPK11, MAPK12 |
| TXK | 7/108 (6.48%) | 5.26x10<sup>-02</sup> | CISH, PIK3R1, STAT1, SOCS1, PTPN6, AKT2, PIK3R3 |
| RASSF1 | 3/25 (12.00%) | 4.18x10<sup>-02</sup> | AKT1, MAP2K6, MAP2K3 |
| RPS3 | 2/17 (11.76%) | 9.71x10<sup>-02</sup> | NFKBIA, NFKB1 |
| RBM4 | 1/75 (1.33%) | 9.07x10<sup>-01</sup> | MAPK3 |
| YES1 | 1/43 (2.33%) | 7.44x10<sup>-01</sup> | PIK3R1 |
| PSMA5 | 2/46 (4.35%) | 4.22x10<sup>-01</sup> | PLK1, USP7 |
| GABPB1 | 2/39 (5.13%) | 3.45x10<sup>-01</sup> | NFKBIA, RELA |
| PTPN22 | 3/62 (4.84%) | 3.05x10<sup>-01</sup> | SOCS1, PTPN6, STAT1 |
| TRIM5 | 1/20 (5.00%) | 4.69x10<sup>-01</sup> | CISH |
| BRD4 | 1/69 (1.45%) | 8.88x10<sup>-01</sup> | MAP2K1 |
| TESK2 | 1/66 (1.52%) | 8.77x10<sup>-01</sup> | STAT1 |

Abbreviations: \*, gene is known to be in the pathway; AS, alternative splicing; LPS, lipopolysaccharide; NEASE, Network-based Enrichment method for AS Events; PBS, phosphate-buffered saline.

## 4 Discussion

LPS, a potent activator of the innate immune system, triggers complex inflammatory responses critical for host defense but is also implicated in various pathologies. Our study identifies diverse levels of associations, ranging from pathways identified to be directly induced by LPS to indirect involvement in immune cell function or cellular homeostasis affected by inflammatory stimuli. This study is the first to perform a characterization of the transcriptomic response of LPS stimulation of human PBMCs using deep sequencing of bulk RNA. The discussion will first briefly review the findings of the global transcriptomic analyses, followed by an in-depth discussion of the findings of the alternative splicing analyses.

### 4.1 Patterns of Global Gene Expression

Briefly, our findings are concordant with other studies that found elevated expression of cytokines associated with LPS stimulation (i.e., IFN-γ, IL-2, IL-4, IL-5, IL-6, IL-8, IL-10, IL-17, and IL-23; Supplementary Figure 2). Pathway impact analysis identified numerous signaling pathways previously identified as responses to LPS stimulation related to modulation (Cytokine-cytokine receptor interaction, RNA Transport, PI3K-Akt, RNA degradation),(60, 61) activation/regulation (Toll-like Receptor, TGF-beta, MAPK, mTOR, NF-kappa B), (60–62) and induction (ErbB, HIF-1).(62, 63)

Two signaling pathways, SNARE Interactions in Vesicular Transport and mRNA Surveillance Pathway, were identified as novel associations with LPS stimulation. SNARE proteins play a crucial role in vesicle trafficking and membrane fusion, which are essential for cellular responses to stimuli like lipopolysaccharides (LPS). SNARE proteins are a family of proteins that form a complex to pull two membranes (like a vesicle and the cell’s plasma membrane) together, allowing them to fuse and release their contents. While LPS stimulation clearly leads to the secretion of cytokines (a process dependent on vesicular transport and therefore, SNAREs), the existing research on this topic has not focused on SNARE proteins as a key regulatory point in the LPS-PBMC response. Given the mediation of vesicles by the SNARE complex for secretion, future research is needed to clarify potential connections in immune cell secretion, phagocytosis, or intracellular trafficking in response to LPS.

Nonsense-mediated decay (NMD), represented by the mRNA Surveillance Pathway, functions to identify and degrade faulty mRNA transcripts that contain premature termination codons (PTCs) and help to regulate the expression levels of many normal, full-length transcripts.(64, 65) In terms of LPS, NMD regulates inflammatory gene expression(66) and the expression of certain pathogen-sensing receptors (e.g., TLRs).(4) While the inflammatory response to LPS is well-studied, the role of NMD in modulating this response is less well known. NMD may play a post-transcriptional regulatory layer that fine-tunes the immune system’s reaction to pathogens.

Our global differential expression results are consistent with previous findings and also identify novel pathways associated with LPS stimulation. For the remainder of the discussion, we focus on the findings of alternative splicing patterns in the immune response to LPS stimulation.

### 4.2 Patterns of Alternative Splicing of the Whole-transcriptome

Concordant with the global gene expression patterns and previous research, pathways involved in Immune system and inflammation, Signaling pathways, Cell adhesion, cytoskeleton and extracellular matrix, Metabolism and biosynthesis, Cellular processes and vesicle trafficking, and Cancer and disease pathways processes were enriched for DAS genes. In terms of the broad inflammatory and immune response, alternative splicing is a widespread regulatory mechanism in many of these pathways. TLRs are discussed in more detail below. Genes involved in the TCR signaling, IL-1 signaling, and Interferon type I signaling pathways all show evidence of alternative splicing that modulates their activity during an immune response. For example, some isoforms of T-cell receptors or cytokine receptors can have altered signaling properties or act as dominant-negative inhibitors. The inflammatory signals from LPS directly influence the activity of these pathways, and the resulting alternative splicing helps to fine-tune the overall immune response, preventing over-activation and tissue damage.

In addition to these known pathways, four pathways that have limited understanding of the response to LPS in the context of alternative splicing include Neddylation, the PI3K/AKT network, Focal Adhesion, and Receptor tyrosine kinases signaling. Through post-transcriptional modification, the Neddylation pathway is a critical regulator of immune and inflammatory responses primarily by controlling the activity of cullin-RING ligases (CRLs).(67–69) Neddylation of cullins activates these ligases, which in turn tag hundreds of proteins for degradation, including many key components of inflammatory signaling pathways. Current evidence for the regulation of this pathway during inflammation is focused on changes in enzyme activity and protein levels,(69) rather than on the production of functionally distinct isoforms through alternative splicing. That said, of the DAS genes identified in the analysis as having effects of AS activity in this pathway, HDAC7(70) and TRIM5(71) have evidence for isoforms with functional relevance to immune responses. While the Neddylation pathway is known to be involved in the LPS response, the role of alternative splicing within this pathway is a new opportunity for research.

The PI3K/AKT network is a major intracellular signaling pathway that controls cell survival, metabolism, and proliferation. It’s a central hub for many signals, including those from TLRs and RTKs, and is a key regulator of the immune response. Although the pathway is well-studied, the impact of alternative splicing on the function of its numerous components (e.g., kinases, phosphatases, and adaptor proteins) in an LPS-induced context is not fully understood. Altered splicing of a negative regulator of this network could lead to its over-activation, contributing to excessive inflammation.

Focal adhesions are dynamic structures that link the cell’s cytoskeleton to the extracellular matrix, playing a vital role in cell migration and cell-to-cell communication. Immune cell migration to sites of inflammation is a hallmark of the LPS response. The proteins that make up focal adhesions, such as integrins and adaptor proteins, are known to have multiple splice variants. However, how LPS stimulation might alter the splicing patterns of these proteins to fine-tune the migration and function of immune cells is largely an unexplored area. Studying this could reveal novel mechanisms by which the body regulates the inflammatory response.

Receptor tyrosine kinases (RTKs) are cell surface receptors that play a crucial role in cell growth, proliferation, and differentiation. While their role in innate immunity is recognized, the full extent to which they are regulated by alternative splicing during an inflammatory response is not well-documented. Many RTKs, such as those for VEGF, EGF, and PDGF, are activated during inflammation. The alternative splicing of these receptors could produce isoforms that lack their extracellular domain, acting as dominant-negative inhibitors, or soluble forms that serve as decoy receptors to reduce signal intensity. This process would act as a crucial negative feedback loop to temper the inflammatory response.

### 4.3 LPS-stimulated Macrophage polarization is associated with alternative splicing

Macrophages are polarized in different conditions according to their environment, with M1 polarization primarily associated with pro-inflammatory responses and M2 polarization with anti-inflammatory responses.(72) Long-term (>12h) stimulation with LPS promotes M1 polarization and decreases M2 polarization.(73) Concordant with these patterns, we see increased levels of M1 Macrophages in LPS-treated cells as compared to PBS-treated cells. No M2 Macrophages were detected in LPS-treated cells, whereas some level was detected in PBS-treated cells. Previous studies evaluating the effects of LPS using scRNAseq have identified two distinct monocyte transcriptional states in response to LPS: an early activation characterized by the expression of chemoattractants and a later pro-inflammatory state characterized by expression of effector molecules.(74) Although we did not see differences associated with LPS treatment in other cell types, future studies should evaluate other deconvolution methods(49) or directly evaluate with single-cell analysis.(74)

In terms of individual genes, CYBB and DHCR7 have been identified as regulators of macrophage polarization.(75) In this study, CYBB was the top SE differentially alternative splicing event upregulated in LPS and was differentially expressed in the global differential expression analysis. In contrast, DHCR7 was not found to be differentially alternatively spliced or differentially expressed in our evaluation. Interestingly, CD163 is a surface marker for a subpopulation of macrophages associated with inflammation.(76) In our study, CD163 is differentially alternatively spliced through A3SS, but not differentially expressed. Given the interest in the development of drug development targeting CD163 in models of inflammation and cancer, (76, 77) an evaluation of the effects of alternative splicing may guide future research. Our findings of differential alternative splicing associated with macrophage differentiation, and recent development in our understanding of the systematic proteomic inflammatory response to LPS stimulation in macrophages,(78) suggest that future research should evaluate the role of differential mRNA splicing in the dysregulation of the inflammatory response.

### 4.4 Alternative splicing in the Toll Like Receptors pathway

Toll like receptors recognize and respond to conserved pathogen-associated molecular patterns (PAMPs)(79) and are well studied as a response to LPS treatment.(80) Concordant with these findings, the Toll-like Receptor (TLR) Signaling pathway (hsa04620) was perturbed in our global transcriptome analysis (Table 2). In addition, several of the globally differentially expressed genes are key players in TLR signaling (TLR5, TLR7, IRAK2, NFKB2, NFKB1A, TLR10) that are directly involved in sensing pathogens, transmitting signals from the cell surface, and triggering the downstream immune response.(81)

Although alternative splicing has been identified as a key aspect of the TLR Signaling pathway response to infection, more research is needed to evaluate this mechanism.(5) One mechanism that terminates persistent TLR signaling as part of the inflammatory response is the alternative splicing of MyD88. Concordant with these hypotheses, in the splicing-aware interaction network enrichment analyses, the Regulation of Toll-like Receptor signaling (WP1449), Toll-like Receptor signaling (WP75), Simplified Depiction of MYD88 Distinct Input-Output Pathway (WP3877), and Toll-like receptor Signaling Related to MyD88 (WP3858) pathways (Tables 5, Figure 6) were enriched for genes with a potential to rewire network interactions due to AS of exons. Furthermore, in the NEASE analysis, TLR4 was identified as the top AS gene (Figure 7) driving the enrichment of the Regulation of Toll-like Receptor Signaling pathway (WP1449) affecting the binding of sixteen other genes by alternative splicing (Table 8), including MyD88.

**Figure 7.**
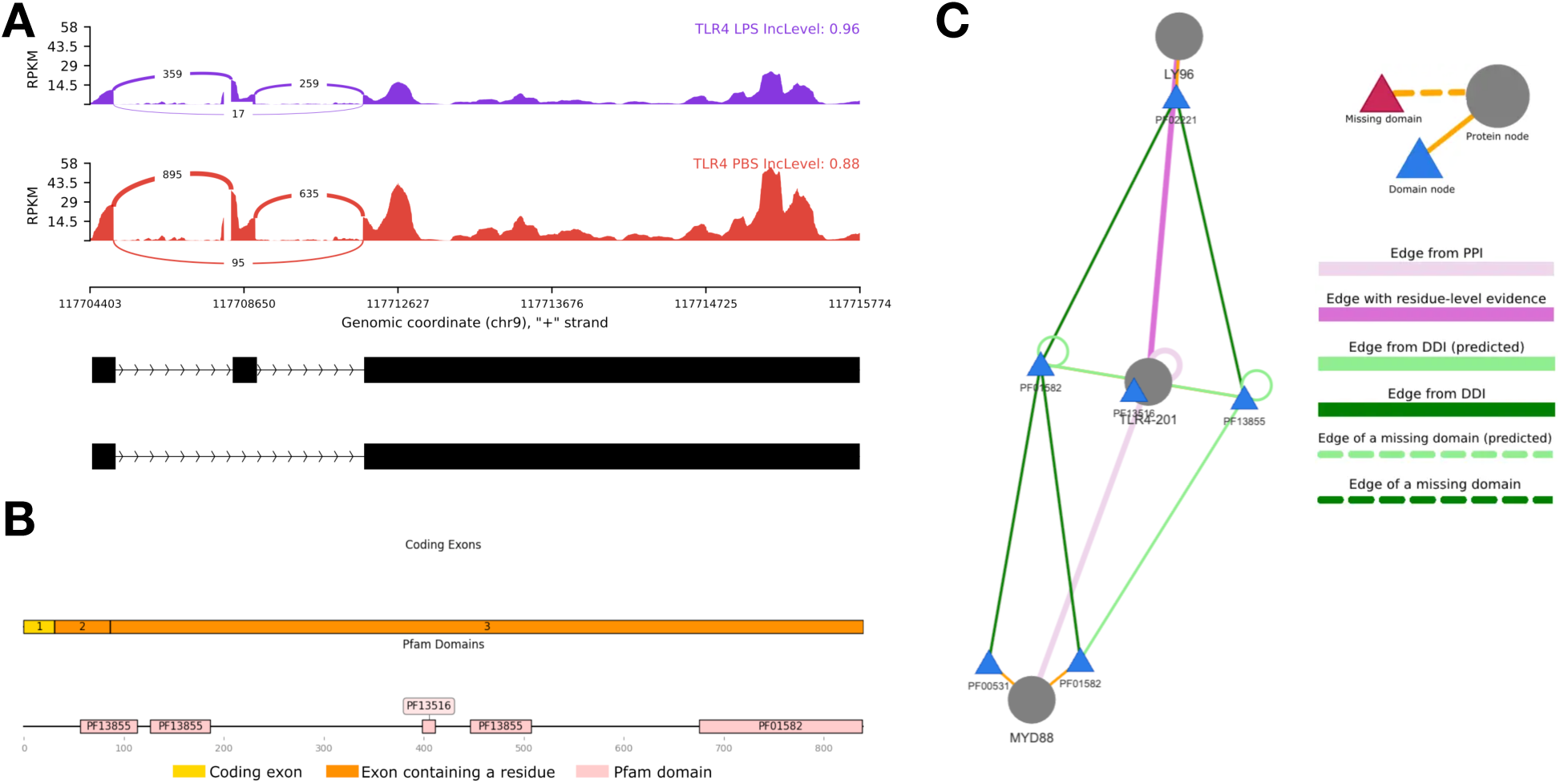
Alternative splicing of TLR4. (A) Sashimi plots showing differential alternative splicing events between the LPS and PBS treatment, involving exon 2 in **TLR4**. The black bars and dashed lines at the bottom represent exons and introns, respectively. Density graphs represent reads per kilobase per million (RPKM) mapped to each genomic position. Arcs represent splice junctions. Numbers represent counts of reads mapped to splice junctions. PSI values are shown on top of the sashimi plots. Numbers and PSI values of events shown in the figure are calculated based on RNA-seq splice junction reads. (B) Domain and exon architecture of the transcript TLR4-201 showing 3 exons and 3 unique protein domains. (C) Visualization of TLR4 isoform-specific interactions of transcripts with their interacting domains with MyD88 and LY96 (MD-2) confirmed by both Protein-Protein Interactions (PPI) and Domain-Domain Interactions (DDI) networks.

In terms of MyD88, a critical adaptor protein in Toll-like receptor signaling, its alternative splicing has been studied mostly in the context of SE events, such as the generation of the MyD88-S isoform, which lacks exon 2 and functions as a dominant-negative regulator of signaling. (82) In this study, the common exon skipping isoforms MyD88-L and MyD88-S were not differentially alternatively expressed. However, differential alternative splicing was observed for an A5SS event with MyD88. While A5SS isoforms of MyD88 have not been widely reported, the presence of other splicing events in the gene suggests that additional isoforms, including those generated by A5SS, may exist but remain underexplored. A5SS events can lead to the inclusion of part of an exon or a shift in the reading frame, potentially producing isoforms with altered protein domains. For MyD88, this may lead to alterations in the Toll/Interleukin-1 Receptor (TIR) domain, which is crucial for receptor interaction, or in the death domain, which is essential for downstream signaling. Such changes might influence the protein’s ability to mediate pro-inflammatory signaling or interact with other signaling molecules.

### 4.5 Alternative splicing of long-noncoding RNAs

There is increasing evidence that long non-coding RNAs (lncRNAs) are an important component of the innate immune response.(83) They are differentially expressed in human monocytes in response to bacterial LPS,(11) and are involved in macrophage polarization.(84) They have a substantial role in the regulation of alternative splicing and, through several mechanisms, can shape the expression of a diverse set of splice isoforms.(85) They are alternatively spliced themselves, resulting in isoform diversity which can affect their stability, localization, and regulatory functions.(86) Two lncRNA genes (MALAT1, PVT1) were identified as DAS in our study that are concordant with previous findings of alternative splicing. MALAT1 is an abundant, conserved, and broadly expressed lncRNA that exhibits an unusual 3′-end processing mechanism.(87) It has been implicated in gene regulations through the modulation of alternative splicing events,(88) cancer prognosis,(89) and autoimmune diseases.(90) MALAT1 is upregulated in lipopolysaccharide (LPS)-activated macrophages, and is hypothesized to function as an autonegative feedback regulator of NF-κB.(91) PVT1 is associated with immune signaling and inflammation, has a role in pretranscription regulation, acts as an epigenetic modulator, and can encode miRNAs.(92) It has received much attention in the past decade for its role in cancer progression(93) and other human diseases.(92) PVT1 exhibits a large number of isoforms which are associated with colorectal cancer tissues,(94) stress,(95) or immune activation (96) and PVT1 splicing at certain splice sites influences target gene expression.(97)

In terms of novel findings, we were unable to identify previous studies identifying alternative splicing events under LPS stimulation or immune-related conditions for much of these DAS lncRNA genes. Two lncRNA genes were identified to have evidence of a role in inflammatory responses (GUSBP11) or gene regulation (JPX), suggesting a strong possibility that alternative splicing could be a regulatory mechanism in this context. GUSBP11 is involved in the broader immune and inflammatory response, including the reaction to LPS stimulation. In a recent study on human gingival fibroblasts,(98) the inhibition of GUSBP11 was found to alleviate LPS-induced cellular inflammation. This effect was mediated through the regulation of microRNA-185-5p (miR-185-5p), indicating a specific molecular mechanism by which GUSBP11 influences the LPS response pathway and a role through alternative splicing (e.g., miRNA sponge (99)). JPX, a lncRNA involved in X chromosome inactivation, targets thousands of genomic sites, preferentially binding promoters of active genes.(100, 101) JPX regulates gene expression through the inhibition of anchor site usage of CCTF and the formation of chromatin loops.(102) Given alternatively spliced *Jpx* RNA acts as a positive regulator of *Xist* in mouse post-X chromosome inactivated cells,(103) alternatively spliced JPX may also play a role in the regulation of gene expression and response to LPS stimulation. Given our sample is exclusively female donors, further research is needed to evaluate the role and molecular regulatory mechanism of alternative splicing of these lncRNAs in response to LPS.

## 5 Limitations and Future Research

Short-read RNA-seq does not directly detect full-length isoforms.(104) In addition, transcriptome-based analyses are unable to evaluate the proteomic phenotype responses.(104) Future studies should utilize long-read RNA-seq and proteome analysis to further characterize alternative splicing due to LPS.(104) This study used the rMATS-turbo tool, which reports differential splicing of canonical alternative splicing events (ASEs). To capture a broader set of AS events, future studies should evaluate for alternative splicing using intron-oriented methods (e.g., LeafCutter,(105) MntJULiP,(106) and MAJIQ(107)). Future research should evaluate ultra-high depth sequencing (150-500M reads) to evaluate lower expressed genes.(59) Future research should evaluate the effects of microRNAs on the regulation of post-transcriptional gene expression of alternatively spliced genes.(108)

Given our findings of common and distinct patterns of global differential expression as compared to differential alternative splicing, alternative splicing offers new insight into the transcriptomic response to LPS stimulation in PBMCs that would be missed through a global transcriptomic evaluation. Through deep sequencing, this study is the first to evaluate for transcriptome-wide patterns of alternative splicing in response to LPS treatment in PBMCs. This study provides a characterization of the transcriptome response in response to LPS treatment in PBMCs and provides potential targets for intervention. Additionally, it offers the opportunity to explore widespread conditions such as sepsis, chronic inflammatory diseases, and other immune-related pathologies.

## Data Availability Statement

The data discussed in this publication have been deposited in NCBI’s Gene Expression Omnibus (Kober et al., 2025) and are accessible through GEO Series accession number GSE308443 (https://www.ncbi.nlm.nih.gov/geo/query/acc.cgi?acc=GSE308443).

## Author Contributions

ECI: Conceptualization, Data curation, Formal analysis, Investigation, Methodology, Project administration, Validation, Visualization, Writing – original draft, Writing – review & editing

AS: Conceptualization, Methodology, Supervision, Writing – review & editing

NT: Writing – review & editing, Validation

JT: Writing – review & editing, Validation

SY: Writing – review & editing, Validation

MS: Writing – review & editing, Validation

AO: Formal analysis, Methodology, Writing – review & editing, Validation, Resources

NWC: Writing – review & editing, Validation

KMK: Conceptualization, Data curation, Data curation, Funding acquisition, Investigation, Methodology, Project administration, Resources, Software, Supervision, Validation, Visualization, Writing – original draft, Writing – review & editing

## Acknowledgments

Esther Chavez-Iglesias is a UCSF PROPEL trainee.

