## Supplemental File 1 for "Alternative Splicing And Global Transcriptome Changes Associated With LPS Stimulation In Human Peripheral Blood Mononuclear Cells"

**This PDF file includes:**

Supplemental Table 1. Demographic and clinical characteristics of the woman donor PBMC samples

Supplemental Table 2. Summary of alternative splicing events

Supplemental Table 3. Enrichment of biological pathways for alternatively spliced genes

Supplemental Table 4. Summary of splice-aware protein-protein network enrichment tests for TLR

Supplemental Figure 1. PCA plots of individual and combined duplicate samples

Supplemental Figure 2. Boxplots of expression for common differentially expressed inflammatory genes

Supplemental Figure 3. Enrichment of the top 10 GO terms for differentially expressed genes

Supplemental Figure 4. Summary of alternatively spliced (AS) genes with affected protein features

**Supplemental Data Tables (in separate documents):**

Supplemental File 2. Global differential gene expression results.

Supplemental File 3. Enrichment of GO categories for global differentially expressed genes results.

Supplemental File 4. Pathway impact analysis of global differential gene expression results.

Supplemental File 5. Differential alternative splicing results.

Supplemental File 6. Pathway enrichment of differential alternative splicing results.

Supplemental File 7. Network enrichment of alternative splicing results.

Supplemental Table 1. Demographic and clinical characteristics of the woman donor PBMC samples.

| Study ID | Age | Ethnicity | Smoker | Weight | Height | Blood Type | Viability | CMV status |
| --- | --- | --- | --- | --- | --- | --- | --- | --- |
| 0035 | 48 | Asian | No | 58 | 166 | O+ | 99.0 | Negative |
| 0036 | 54 | Caucasian | No | 86 | 180 | A+ | 99.3 | Negative |
| 0037 | 50 | Black | No | 100 | 155 | O+ | 98.0 | Unknown |

Abbreviations: CMV, Cytomegalovirus

Supplemental Table 2 - Summary of alternative splicing events.

| Event Type | | Total Events JC | | Total Events JCEC | | Sig Events JC | Sig Events JC Sample Higher Inclusion | | Sig Events JC Sample 2 Higher Inclusion | | Sig Events JCEC | | Sig Events JCEC Sample Higher Inclusion | Sig Events JCEC Sample 2 Higher Inclusion |
| --- | --- | --- | --- | --- | --- | --- | --- | --- | --- | --- | --- | --- | --- | --- |
| SE | | 158,340 | | 160,586 | | 5,183 | 2,467 | | 2,716 | | 5,400 | | 2,498 | 2,902 |
| MXE | | 49,456 | | 50,108 | | 1,700 | 851 | | 849 | | 1,684 | | 804 | 880 |
| RI | | 22,862 | | 22,917 | | 1,203 | 434 | | 769 | | 1,294 | | 476 | 818 |
| A3SS | | 67,759 | | 67,861 | | 1,209 | 588 | | 621 | | 1,205 | | 595 | 610 |
| A5SS | | 39,622 | | 39,722 | | 737 | 372 | | 365 | | 786 | | 397 | 389 |

Abbreviations: A3SS, alternative 3’ splice sites; A5SS, alternative 5’ splice sites; AS, alternatively spliced; MXE, mutually exclusive exons; RI, retained introns; SE, spliced exon; Sig, significant; JC, junction counts; JCEC, junction counts and exon counts.

Supplemental Table 3. Enrichment of biological pathways for alternatively spliced genes across event types between LPS and PBS-treated PBMCs from three women donors.

| Event Type | Database | AS Genes  Tested (n) | Pathways Tested  (n) | Pathways Enriched  (n) |
| --- | --- | --- | --- | --- |
| SE | WikiPathway | 940 | 446 | 58 |
|  | Elsevier |  | 1,190 | 132 |
|  | BioPlanet |  | 1,071 | 139 |
| MXE | WikiPathway | 240 | 214 | 4 |
|  | Elsevier |  | 513 | 14 |
|  | BioPlanet |  | 503 | 20 |
| RI | WikiPathway | 356 | 376 | 134 |
|  | Elsevier |  | 1,035 | 489 |
|  | BioPlanet |  | 808 | 237 |
| A3SS | WikiPathway | 223 | 248 | 1 |
|  | Elsevier |  | 580 | 4 |
|  | BioPlanet |  | 543 | 5 |
| A5SS | WikiPathway | 146 | 199 | 22 |
|  | Elsevier |  | 552 | 97 |
|  | BioPlanet |  | 402 | 35 |

Abbreviations: A3SS, alternative 3’ splice sites; A5SS, alternative 5’ splice sites; AS, alternatively spliced; MXE, mutually exclusive exons; RI, retained introns; SE, spliced exon.

Supplemental Table 4. Summary of splice-aware protein-protein network enrichment tests for Toll Like Receptor, TLR4, and MyD88 related pathways from the Reactome (R-HSA) and WikiPathways (WP) databases associate with differentially alternatively spliced genes of human PBMCs treated with LPS or PBS predicted to impact protein-protein network interactions.

| Pathway ID | Pathway name | AS genes (SE events only)  (number of interactions affecting the pathway) | FDR | NEASE score |
| --- | --- | --- | --- | --- |
| WP1449 | Regulation of toll-like receptor signaling pathway | BRD4 (1), RASSF1 (3), RBM4 (1), GRK3 (3), PTPN22 (3), RPS3 (2), NFKBIE (9), IL1R1 (4), TESK2 (1), MAP2K2 (17), ANKRD28 (8), ACVR1B (8), PSMA5 (2), TLR4 (16), GABPB1 (2), YES1 (1), TXK (7), PIK3C2A (4), TRIM5 (1) | 4.04x10^-6^ | 23.74 |
| WP75 | Toll-like Receptor Signaling Pathway | BRD4 (1), RASSF1 (3), RBM4 (1), GRK3 (3), PTPN22 (1), RPS3 (2), NFKBIE (7), IL1R1 (3), TESK2 (1), MAP2K2 (15), ANKRD28 (5), ACVR1B (6), TLR4 (12), GABPB1 (2), YES1 (1), TXK (4), PIK3C2A (4) | 1.39x10^-4^ | 17.69 |
| WP3877 | Simplified Depiction of MYD88 Distinct Input-Output Pathway | RPS3 (1), RNF185 (1), NFKBIE (3), IL1R1 (1), MAP2K2 (1), ANKRD28 (1), TLR4 (7), UBE2B (2) | 1.08x10^-3^ | 9.38 |
| WP3858 | Toll-like receptor signaling related to MyD88 | RPS3 (1), NFKBIE (7), IL1R1 (1), MAP2K2 (2), ANKRD28 (4), ACVR1B (1), TLR4 (9), GABPB1 (2) | 4.81x10^-3^ | 7.63 |

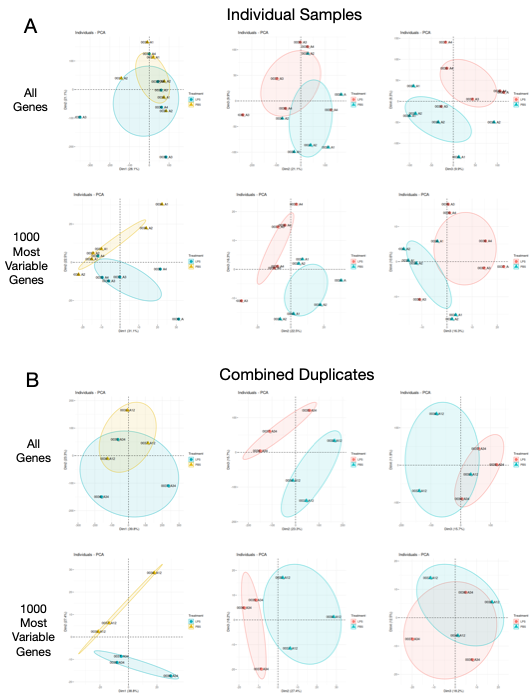

Supplemental Figure 1. Plots of the first four dimensions of a principal components analysis (PCA) of gene expression levels for peripheral blood mononuclear cells (PBMCs) from three donors treated with either lipopolysaccharide (LPS) or Phosphate-buffered saline (PBS) for (A) all individual samples and (B) combined duplicates using all genes or the 1000 most variable genes.

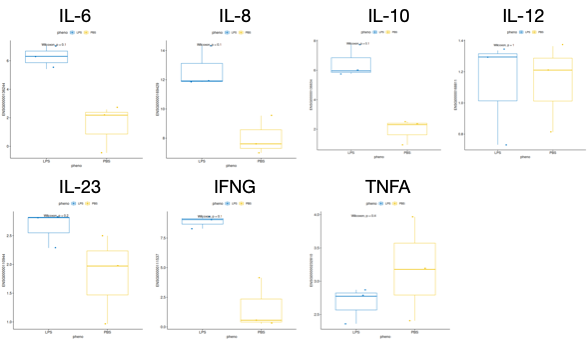

Supplemental Figure 2. Boxplots of expression for LPS and PBS-treated human PBMCs for pro- and anti-inflammatory genes commonly associated with immune response and LPS treatment.

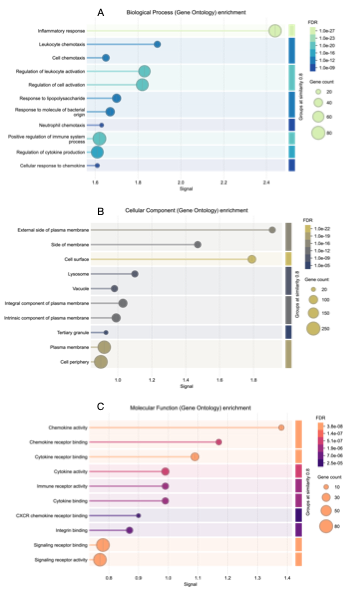

Supplemental Figure 3. Enrichment of the top 10 GO terms for genes differentially expressed between LPS and PBS.

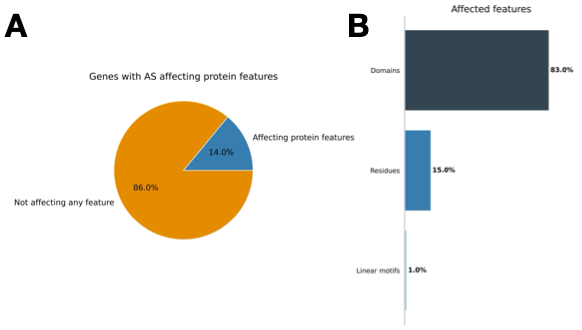

Supplemental Figure 4. Summary of alternatively spliced (AS) genes with affected protein features associated with LPS or PBS treatment of human PBMCs. (A) Gene affecting protein features. (B) Summary of affected features.
